# Blue turns to grey - Palaeogenomic insights into the evolutionary history and extinction of the blue antelope (*Hippotragus leucophaeus*)

**DOI:** 10.1101/2022.04.12.487785

**Authors:** Elisabeth Hempel, Faysal Bibi, J. Tyler Faith, Klaus-Peter Koepfli, Achim M. Klittich, David A. Duchêne, James S. Brink, Daniela C. Kalthoff, Love Dalén, Michael Hofreiter, Michael V. Westbury

**Author notes:** deceased (James S. Brink). shared senior authorship (Michael Hofreiter, Michael V. Westbury). Corresponding author: Elisabeth Hempel.

## Abstract

The blue antelope (*Hippotragus leucophaeus*) is the only large African mammal species to have become extinct in historical times, yet no nuclear genomic information is available for this species. A recent study showed that many alleged blue antelope museum specimens are either roan (*H. equinus*) or sable (*H. niger*) antelopes, further reducing the possibilities for obtaining genomic information for this extinct species. While the blue antelope has a rich fossil record from South Africa, climatic conditions in the region are unfavourable to the preservation of ancient DNA. Nevertheless, we recovered two blue antelope draft genomes, one at 3.4x mean coverage from a historical specimen (~200 years old) and one at 2.1x mean coverage from a fossil specimen dating to 9,800–9,300 cal BP, making it currently the oldest palaeogenome from Africa. Phylogenomics show that blue and sable antelope are sister species, confirming previous mitogenomic results, and demonstrate ancient gene flow from roan into blue antelope. We show that blue antelope genomic diversity was much lower than in roan and sable antelopes, indicative of a low population size since at least the early Holocene. This supports observations from the fossil record documenting major decreases in the abundance of blue antelope after the Pleistocene-Holocene transition. Finally, the persistence of this species throughout the Holocene despite low population size suggests that colonial-era human impact was likely a decisive factor in the blue antelope’s extinction.

## Introduction

Earth is currently undergoing a major biodiversity crisis that is a direct result of human activities (Ceballos et al. 2015). The consequences for many species include considerable reductions in population size and associated losses of genetic diversity that can lead to reduced fitness and adaptability. Today, through the continuing development of next generation sequencing technologies and advancement of ancient DNA (aDNA) techniques, it is feasible to retrieve genomic information from extinct and non-model organism species (Westbury et al. 2020; Barlow et al. 2021; Sánchez-Barreiro et al. 2021), thereby facilitating inferences about threats facing many present-day species. One especially powerful use of these genomes is to reconstruct the evolutionary history of species in relation to environmental change and human impacts. In this regard, Africa is hugely understudied, likely due to its environmental conditions, with high temperatures being detrimental to DNA preservation (Smith et al. 2001; Bollongino et al. 2008; Hofreiter et al. 2015). To date, most aDNA studies on African samples have been restricted to DNA enrichment approaches and focused on human remains (Skoglund et al. 2017; Vicente and Schlebusch 2020; Lipson et al. 2022), with very few investigating other fossil fauna (e.g., Mathieson et al. 2020).

One species that likely fell victim to human impacts during the beginning of the current biodiversity crisis is the blue antelope, *Hippotragus leucophaeus* (Pallas, 1766). The blue antelope belongs to the bovid tribe Hippotragini, which comprises the extant genera *Hippotragus*, *Oryx* and *Addax*. It went extinct ~1800 AD (Lichtenstein 1811, 1814) and therefore represents the first – and so far only – historical extinction of a large African mammal species (Harper 1945), though several subspecies have become extinct in the last few hundred years (e.g., quagga (*Equus quagga quagga*), bubal hartebeest (*Alcelaphus buselaphus buselaphus*), cape warthog (*Phacochoerus aethiopicus aethiopicus*), (d’Huart and Grubb 2001; Hack et al. 2008; IUCN SSC Antelope Specialist Group 2017). The blue antelope had a distinct white patch in front of the eyes (Pallas 1767), and its pelt was perceived as bluish while alive (Lichtenstein 1814; von Schreber and Goldfuß 1836) – perhaps similar to the blue wildebeest (*Connochaetes taurinus*) or the nilgai (*Boselaphus tragocamelus*) (Mohr 1967) – turning greyish post-mortem (Lichtenstein 1814; von Schreber and Goldfuß 1836) (fig. 1B). Like its extant relatives, the blue antelope was a grazer (Mohr 1967; Klein 1974, 1987). Historically, the blue antelope was endemic to a very small area (~4,300 km^2^) between Swellendam, Caledon, and Bredasdorp in South Africa (Skead 1980; Kerley et al. 2009). However, its Quaternary fossil record demonstrates a broader prehistoric range (fig. 1A), with fossil occurrences throughout the Cape Floristic Region and extending into the highlands of Lesotho (Klein 1974; Plug 1997; Faith and Thompson 2013; Avery 2019). Blue antelopes are particularly ubiquitous and abundant in Pleistocene archaeological and palaeontological assemblages from South Africa’s Western Cape Province, contracting in range and abundance at the onset of the Holocene (Faith 2011). Its occurrence in both late Pleistocene and Holocene archaeological sites implies a long history of human predation on blue antelope that spans at least the past ~100,000 years (e.g., Klein 1976; Faith 2013), with evidence from Elandsfontein indicating that hominins and blue antelope have overlapped since the mid-Pleistocene (~1 Ma to 600 ka; Klein et al. 2007). The blue antelope’s extinction is hypothesised to have been the result of several anthropogenic drivers, including landscape transformation (Faith and Thompson 2013), overhunting by European colonists (FitzSimons 1920; Harper 1945), competition with and habitat deterioration by livestock (Klein 1974, 1987), and disruption of migratory pathways in prehistoric and colonial times (Faith and Thompson 2013). Additionally, the potentially detrimental role of stochastic processes on a small population size has also been considered (Kerley et al. 2009). Unfortunately, due to its early demise, our primary sources of knowledge of the blue antelope’s ecology and evolutionary history has so far been limited to information from the fossil record (Klein 1974, 1987; Faith 2011; Faith and Thompson 2013) and historical mitochondrial data (Robinson et al. 1996; Themudo and Campos 2018; Hempel et al. 2021a).

**Fig. 1.**
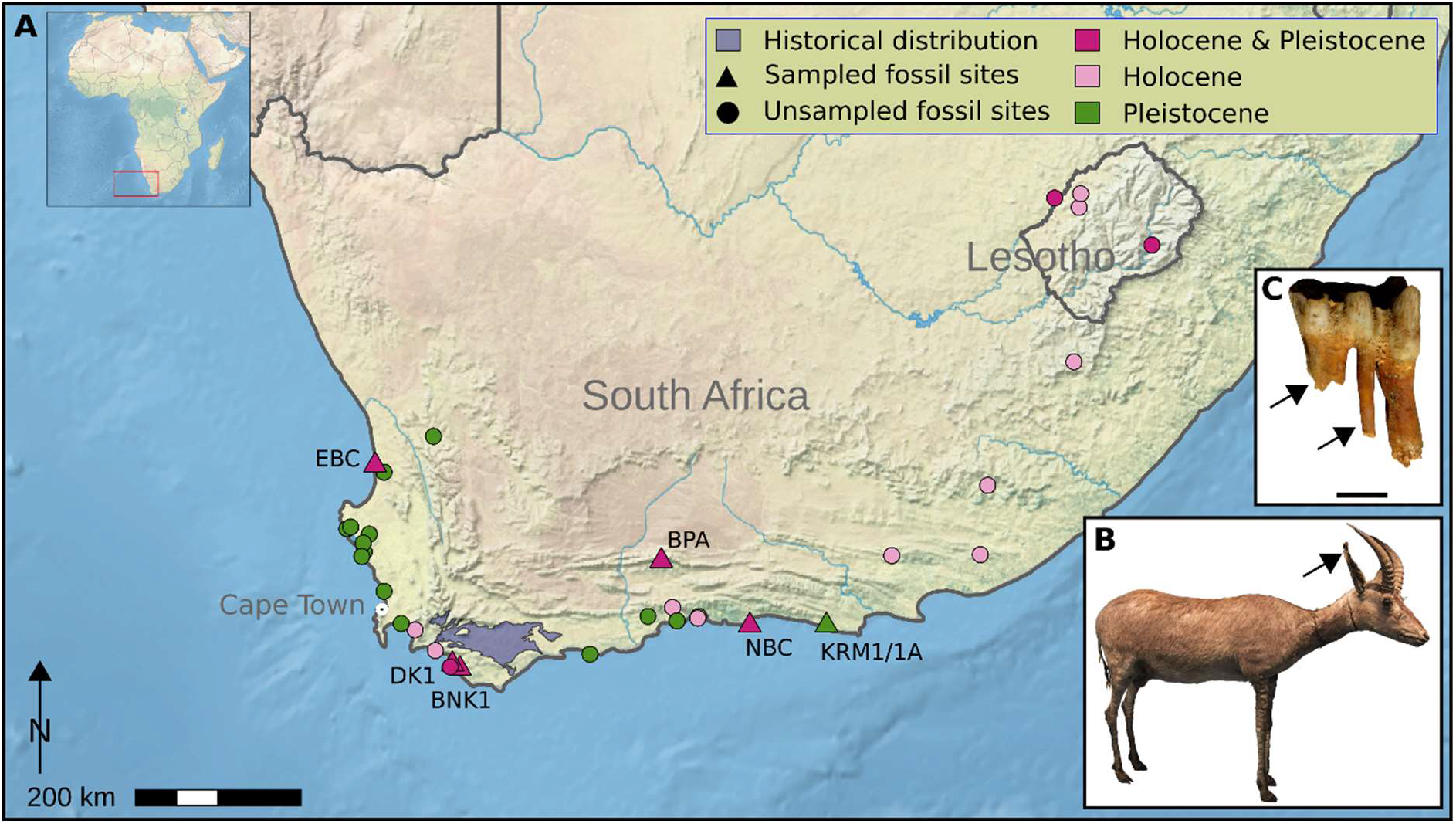
Blue antelope distribution and specimens. (**A**) Historical distribution (light purple at the southern tip) and Quaternary fossil occurrences of blue antelope (Skead 1980; Kerley et al. 2009; modified from Avery 2019). Fossil sites: EBC = Elands Bay Cave, DK1 = Die Kelders Cave 1, BNK1 = Byneskranskop 1, BPA = Boomplaas Cave, NBC = Nelson Bay Cave, KRM1/1A = Klasies River Mouth 1/1A (base map: https://www.naturalearthdata.com, prepared in QGIS v2.18 (https://qgis.org)). (**B**) Historical mounted skin of a young male blue antelope (*Hippotragus leucophaeus*) from the Swedish Museum of Natural History, Stockholm, Sweden (NRM 590107); arrow indicates area sampled for aDNA extraction (**C**) Fossil lower left deciduous fourth premolar (NBC RB4 D3) of a blue antelope from Nelson Bay Cave (curated at Iziko Museums of South Africa, Cape Town, South Africa); scale is one centimetre; arrows indicate areas sampled for aDNA extraction. Photo credits: NBC RB4 D3: J.T. Faith, courtesy: Archaeology Unit, Iziko Museums of South Africa; NRM 590107: Hempel et al. 2021a.

The relationships among the three *Hippotragus* species, the roan (*H. equinus*), the sable (*H. niger*) and the blue antelope, have so far only been investigated using morphological evidence (Vrba and Gatesy 1994) and mitochondrial sequences (Robinson et al. 1996; Themudo and Campos 2018; Hempel et al. 2021a), leaving their phylogeny unresolved at the nuclear genome level. Considering that mitochondrial DNA data can present an incomplete picture of the evolutionary history of a species and can be confounded by past gene flow events (e.g., Larsen et al. 2010; Reich et al. 2010; Edwards et al. 2011; Hailer et al. 2012; Westbury et al. 2020; Liu et al. 2021), nuclear data is necessary to conclusively resolve phylogenetic relationships.

Here, we present the nuclear genome of a ~200 year old blue antelope specimen as well as a palaeogenome from an early Holocene fossil specimen (~9,300–9,800 years BP) recovered from Klein’s 1971–1972 excavation of archaeological deposits at Nelson Bay Cave, located on the Robberg Peninsula at Plettenberg Bay (Klein 1972; see Materials and Methods for further details). To our knowledge, the palaeogenome extracted from this early Holocene fossil specimen (NBC RB4 D3) is currently the oldest palaeogenome from Africa. Using these genomes, we investigate and date the phylogenomic relationships of the blue antelope and show that the blue antelope was more closely related to the sable than to the roan antelope, but with detectable past gene flow between the roan and the blue antelope. Furthermore, we demonstrate that the blue antelope had very low diversity compared to its congeners, likely since at least the beginning of the Holocene.

## Results

### Sequencing results

Using shotgun sequencing, we obtained both a nuclear and a mitochondrial genome from a fossil specimen from Nelson Bay Cave (NBC RB4 D3) (Klein 1972, 1983) dating to 9,300–9,800 cal BP (Loftus et al. 2016) at 2.14x and 234.67x mean coverages, respectively. In addition, we generated a nuclear genome of the ~200 year old historical specimen NRM 590107 from the Swedish Museum of Natural History, Stockholm, Sweden, with a mean coverage of 3.44x. Details on read numbers and mapping statistics can be found in supplementary tables S2 and S3. MapDamage v2.2.0 (Jónsson et al. 2013) analysis of DNA sequences obtained from the NBC RB4 D3 specimen showed typical patterns of damage (caused by cytosine deamination) for an aDNA sample of its age (supplementary figs. S1A and S1B). The same was true for NRM 590107, but in addition, this sample also showed an elevated level of guanine to thymine transversions (supplementary figs. S2A and S2B). This pattern could originate from a hydrogen peroxide treatment of the sample or any other treatment causing oxidative damage, since oxidative DNA damage results in 8-hydroxy-guanine (Kvam and Tyrrell 1997; Nohmi et al. 2005) and this modified base pairs with adenine, resulting in guanidine to thymine changes. However, such a treatment has not been documented for the individual from the Swedish Museum of Natural History.

### Phylogenetic relationships

Nuclear genome phylogenetic analyses using neighbour-joining and 20 kb sliding window trees under the multi-species coalescent supported the monophyly of all three *Hippotragus* species and showed that the blue antelope is more closely related to the sable than to the roan antelope (fig. 2, supplementary fig. S3). Out of the 20 kb sliding window trees, 50.46% were concordant with the species tree (supplementary table S9). Bayesian molecular dating using MCMCtree yielded a median posterior age of the split between the blue and sable antelopes of 1.67 Mya, with a 95% credibility interval ranging between 1.48–1.86 Mya. The split between the roan and sable/blue antelopes had a median of 2.86 Mya and a 95% credibility interval ranging between 2.55–3.19 Mya (fig. 2).

**Fig. 2.**
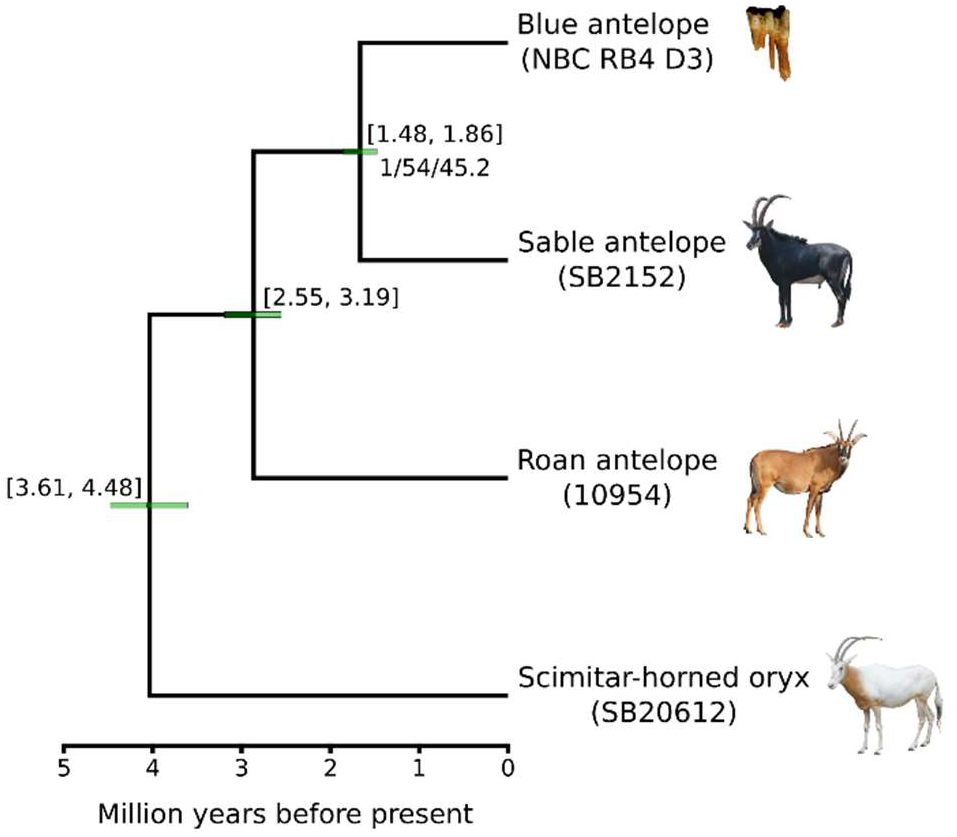
Phylogenetic analysis. Dated phylogenetic tree computed using nuclear and mitochondrial data from the three *Hippotragus* species. The tree topology was inferred using a data set that excluded transition-only sites, under the multi-species coalescent model using ASTRAL v4.10 (Rabiee et al. 2019), while the timescale was inferred using MCMCtree v4.9 (dos Reis and Yang 2011). Node annotations above age credibility interval bars show the credibility intervals range in Mya, while the node label below the bar shows branch support values (local posterior probability, gene concordance factor, site concordance factor). As outgroup the scimitar-horned oryx was used (Humble et al. 2020). Roan and sable antelope raw data are from Gonçalves et al. (2021) and Koepfli et al. (2019). Photo credits: NBC RB4 D3: J.T. Faith, courtesy: Archaeology Unit, Iziko Museums of South Africa; NRM 590107: Hempel et al. 2021a; roan antelope: Charles J. Sharp, wikimedia commons, CC-BY 4.0; sable antelope: Paulmaz, wikimedia commons, CC-BY 3.0; scimitar-horned oryx: E. Hempel.

The mitochondrial genome maximum-likelihood phylogeny was congruent with the nuclear phylogeny (fig. 2, supplementary figs. S3 and S4A). The mitochondrial phylogeny using transversions only (binary format) also yielded an identical topology with respect to the grouping of the three species (supplementary fig. S4B).

### Gene flow

Both blue antelope individuals showed significantly higher levels of shared derived alleles with the roan antelope compared to the sable antelope, suggestive of gene flow between blue and roan antelope (fig. 3A, supplementary table S7). Sliding window phylogenetic tree analyses also suggested gene flow between blue and roan antelope, as more windows contained phylogenies with a closer affinity between roan and blue antelope than between roan and sable antelope (fig. 3C, supplementary table S8 and S9). Moreover, by comparing multiple individuals per species, the sliding window analysis showed that gene flow between roan and blue antelope occurred after the split of blue and sable antelope but before the split of the most recent common ancestors of all roan and also the two blue antelope individuals analysed in our study.

**Fig. 3.**
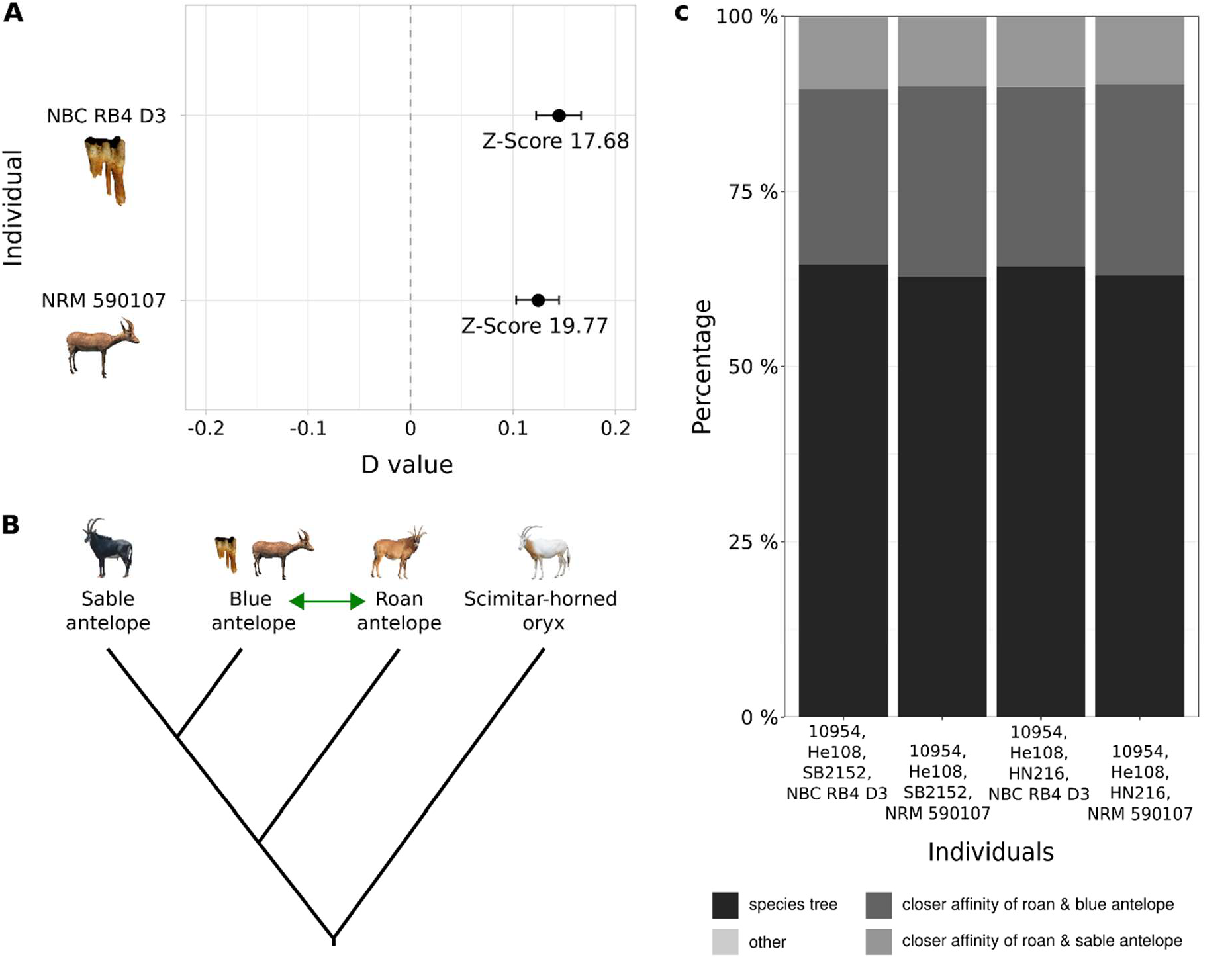
Gene flow analyses for the three *Hippotragus* species. (**A**) *D* values and Z-scores of *D* statistics for the fossil (NBC RB4 D3) and historical (NRM 590107) blue antelope specimens showing significant gene flow between blue and roan antelope; analysis performed with Dstat (transversions only version, https://github.com/jacahill/Admixture). (**B**) Tested topology for *D* statistics according to the 20 kb sliding window multi-species coalescent and the neighbour-joining phylogenies (fig. 2 and supplementary fig. S3) with blue antelope in position two. The green double-arrow indicates gene flow between blue and roan antelope with unknown directionality. (**C**) Percentages of tree topologies in the 100 kb sliding window tree analysis. In every combination more windows show a closer affinity between roan and blue antelope than between roan and sable antelope. Topologies in category “other” were so rare that they are not visible in the figure. Photo credits: see fig. 2.

To determine the directionality of the gene flow, we utilised the branch lengths between roan and sable antelope in window trees where the roan and blue antelope form sister lineages. It was expected that in case of gene flow from roan into blue antelope the branch lengths between roan and sable antelope would be in agreement with the species tree topology, whereas gene flow from blue into roan antelope would lead to a higher similarity between roan and sable antelope and thereby to a shorter branch length compared to the species tree topology (fig. 4A). The sliding window analysis with branch length calculations showed that the branch lengths of roan and sable antelope were similar for the inferred species tree topology and the alternative topology placing blue and roan as sister lineages. Assuming that the most frequently found topology is the species tree topology, the next most frequent topology presumably results from both incomplete lineage sorting and/or gene flow. A bimodal distribution would have been expected if there had been any occurrence of gene flow from the blue into the roan antelope. One mode would reflect the relatively ancient species divergence between the roan and the sable, while the other mode would reflect the more recent divergence between the introgressed blue antelope loci and the sable. Since the distributions of branch lengths between roan and sable were unimodal and approximately equal between windows resulting in the species tree topology and the second most frequent topology, the direction of gene flow was most likely from the roan into the blue antelope (fig. 4B).

**Fig. 4.**
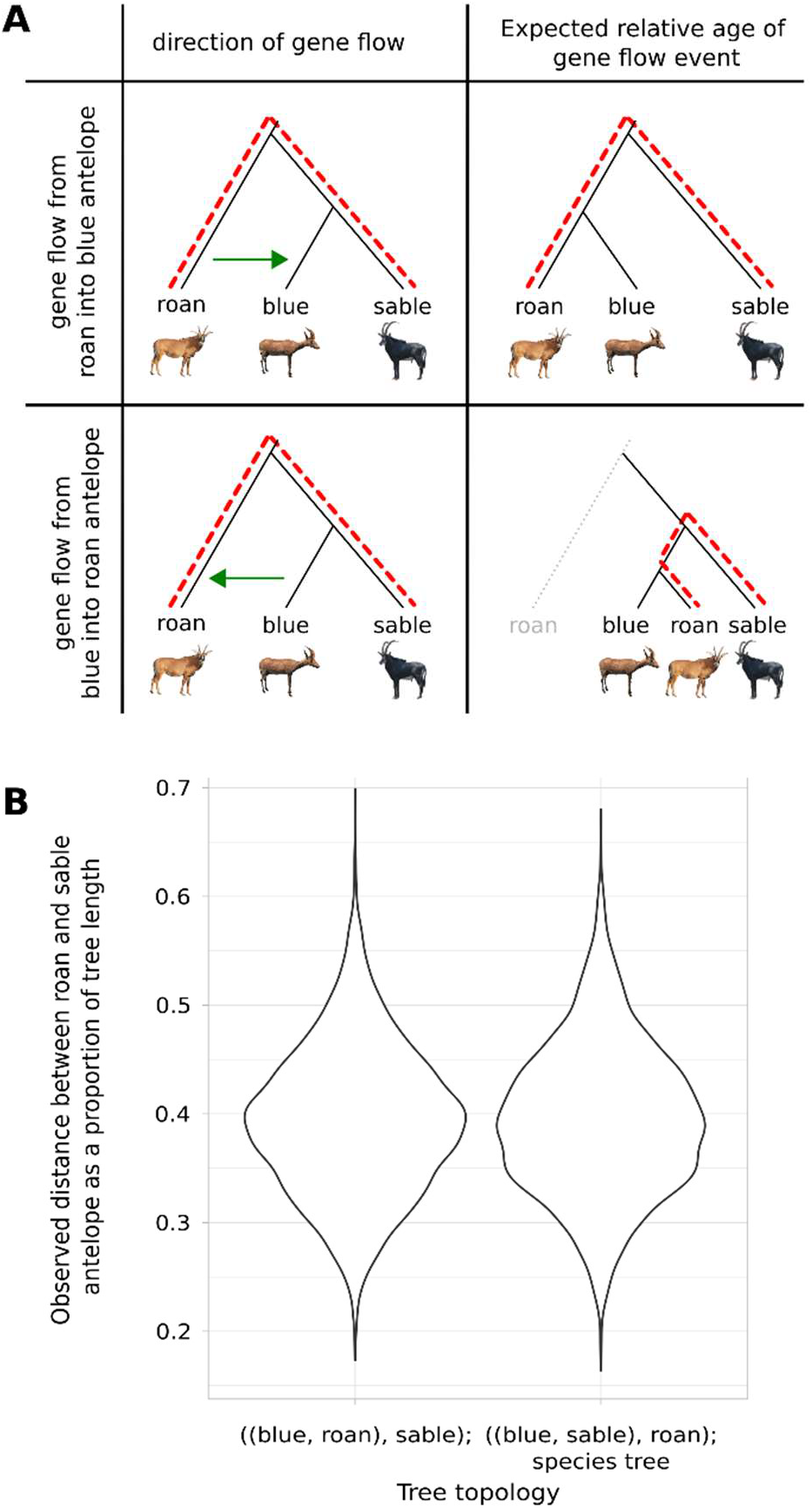
Gene flow direction analysis. (**A**) Expected outcome of changes in genetic distances for gene flow (green arrow) from roan into blue antelope and blue into roan antelope. Dashed red lines illustrate the expected distances between roan and sable antelope. (**B**) Genetic distances between roan and sable antelope as a proportion of the tree length performed with WindowTrees v1.0.0 (https://github.com/achimklittich/WindowTrees). Photo credits: see fig. 2.

### Nuclear and mitochondrial diversity

Calling heterozygous positions to estimate the nuclear genomic diversity of the two low coverage genomes was not a reliable option. Therefore, we estimated diversity through an alternative measure. By uniquely evaluating each substitution type, we confirmed elevated cytosine to thymine and guanine to thymine levels found via mapDamage (supplementary figs. S1 and S2). After removing substitutions that could have been caused by DNA damage, our results show that pairwise differences are related to mean heterozygosity in roan and sable antelope. Therefore, we concluded that pairwise differences could be used to reliably estimate species-wide nuclear diversity even when using low coverage genomes (fig. 5, supplementary tables S10–12 and S16).

**Fig. 5.**
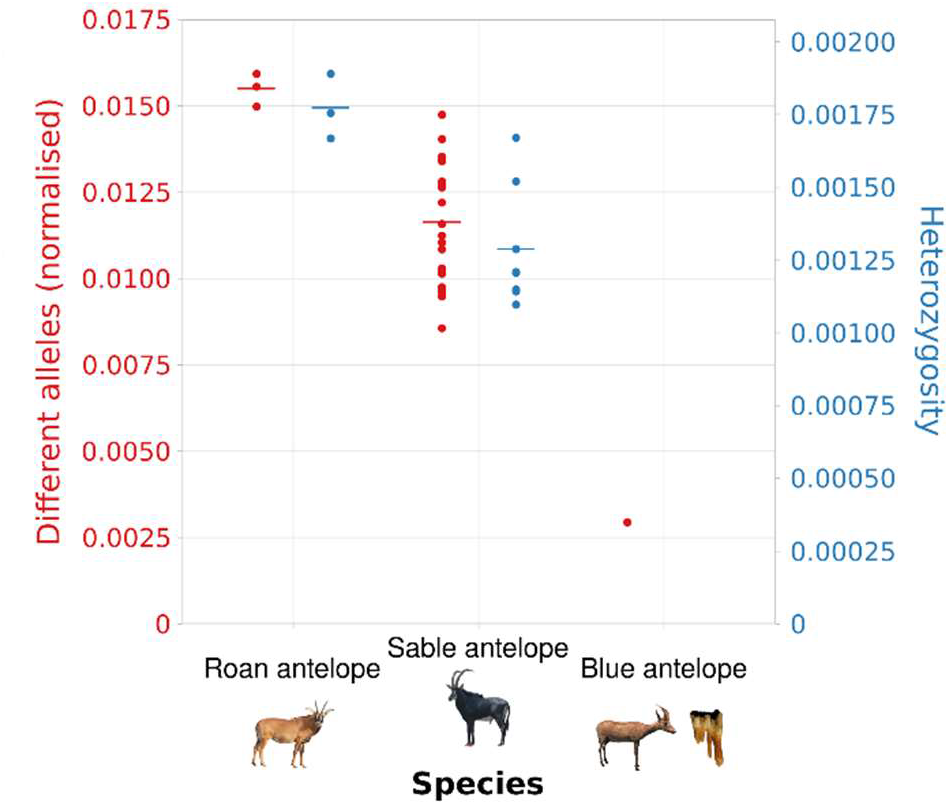
Nuclear diversity analysis. Species-wide nuclear diversity of the three *Hippotragus* species computed from pairwise comparisons (data normalised by dividing by the number of sites used in the analysis - 2,243,953 sites; genomes for heterozygosity subsampled to 4.26x mean coverage, the lowest mean coverage in the roan/sable antelope data set). Each dot represents a pairwise comparison. Lines represent means. Heterozygosity was not estimated for the blue antelope due to low coverage and ancient DNA damage. Photo credits: see fig. 2.

Pairwise difference estimates showed that the blue antelope had much lower nuclear diversity than the roan and sable antelopes (fig. 5). The roan antelope showed the highest nuclear diversity among the three *Hippotragus* species, with the sable antelope showing values that were lower than the roan antelopes’ but still much higher than those of the blue antelope. Comparing subsampled and not subsampled input data for the pairwise difference estimates showed that coverage can have an influence, but not to the extent that it could result in the substantially reduced nuclear diversity of the blue antelope observed relative to the other species (supplementary fig. S5, supplementary tables S13–15).

To estimate the mitochondrial genome diversity of the blue antelope, we generated two haplotype networks, one using complete and the other using partial mitochondrial genomes. The removal of all ambiguities/missing data resulted in an alignment length of 16,492 bp for the complete mitochondrial alignment and 6,300 bp for the partial mitochondrial alignment. We found 15 segregating sites and a nucleotide diversity of 0.00061 in the alignment with complete mitochondrial genomes (supplementary fig. S6A), and 10 segregating sites and a very similar nucleotide diversity of 0.00067 for the alignment with partial mitochondrial genomes (supplementary fig. S6B).

## Discussion

Using ancient DNA (aDNA) techniques, we sequenced the first nuclear genomes of the blue antelope to study its evolutionary history and genetic diversity, and to provide insights into the only historical extinction of a large African mammal species to date. The retrieval of aDNA from African fossils is a challenging task because of prevailing environmental conditions, namely high temperatures, which facilitate DNA degradation (Smith et al. 2001; Bollongino et al. 2008; Hofreiter et al. 2015). For this reason, few studies have succeeded in recovering aDNA from Africa, and the few exceptions were mainly focused on humans and used genomic enrichment approaches (Vicente and Schlebusch 2020; Lipson et al. 2022). Nonetheless, despite such challenges, we were able to successfully sequence a sample that, with an age of 9,300–9’800 years BP, currently represents the oldest palaeogenome from Africa. Our success demonstrates that palaeogenomes can be recovered successfully from southern African sites, setting the stage for future genomic studies from this area. It is worth noting that our fossil specimen originates from cave deposits in South Africa’s southern Cape region, which should have higher potential for aDNA recovery than, for example, fossils from fluvial sediments in tropical Africa (Hofreiter et al. 2015).

Our finding that the blue antelope is more closely related to the sable than to the roan antelope on a nuclear level confirms results from studies based on mitochondrial genome sequences (Robinson et al. 1996; Themudo and Campos 2018; Hempel et al. 2021a) and contradicts phylogenies relying on morphological evidence, which placed roan and blue antelope as sister species (Vrba and Gatesy 1994). Since mitochondrial DNA represents a single genetic locus, it was not certain if the mitochondrial phylogeny alone could reliably settle the question of the relationships among the three *Hippotragus* species, since it could have been biased by introgression or incomplete lineage sorting as has been shown in other species (e.g., Barlow et al. 2018; Rakotoarivelo et al. 2019; Westbury et al. 2020). However, our phylogenomic results confirm the relationships among the recent species of *Hippotragus* on the nuclear level, although they place the split of roan and sable/blue antelope as well as the split between sable and blue antelope younger than these have been dated from mitochondrial genomes (Themudo and Campos 2018). The estimated divergence age of ~1.67 Mya between the sable and blue antelope strongly suggests that the blue antelope represents a separate species, which resolves the long-going discussion on whether the blue antelope was a distinct species or not (Smith 1849; Kohl 1886; Mohr 1967).

Both blue antelope specimens showed significant gene flow with the roan antelope relative to the genomic relationships between the roan and the sable antelope. This may look surprising at first, since the extant and historical populations of the roan and the sable antelope overlap considerably. However, the ranges of the roan and the blue antelope also overlapped in the southern Cape region during the late Pleistocene and Holocene (fig. 6; Klein 1972; Faith 2012; Avery 2019). Interspecific gene flow is known to be more likely when population size is low since the chances of finding a conspecific mating partner are reduced (Hubbs principle) (Hubbs 1955; Jansson et al. 2007; Crossman et al. 2016; Vaz Pinto et al. 2016). Although we were unable to directly date the gene flow event(s), we could show that gene flow occurred after the split of blue and sable antelope but before the last common ancestors of the roan antelope individuals included in this study. While we did not detect evidence for more recent interspecific gene flow, it may have been biologically possible since roan and blue antelope have the same evolutionary distance as roan and sable antelope, which are likely still able to produce viable offspring (Robinson and Harley 1995; Vaz Pinto et al. 2016).

**Fig. 6.**
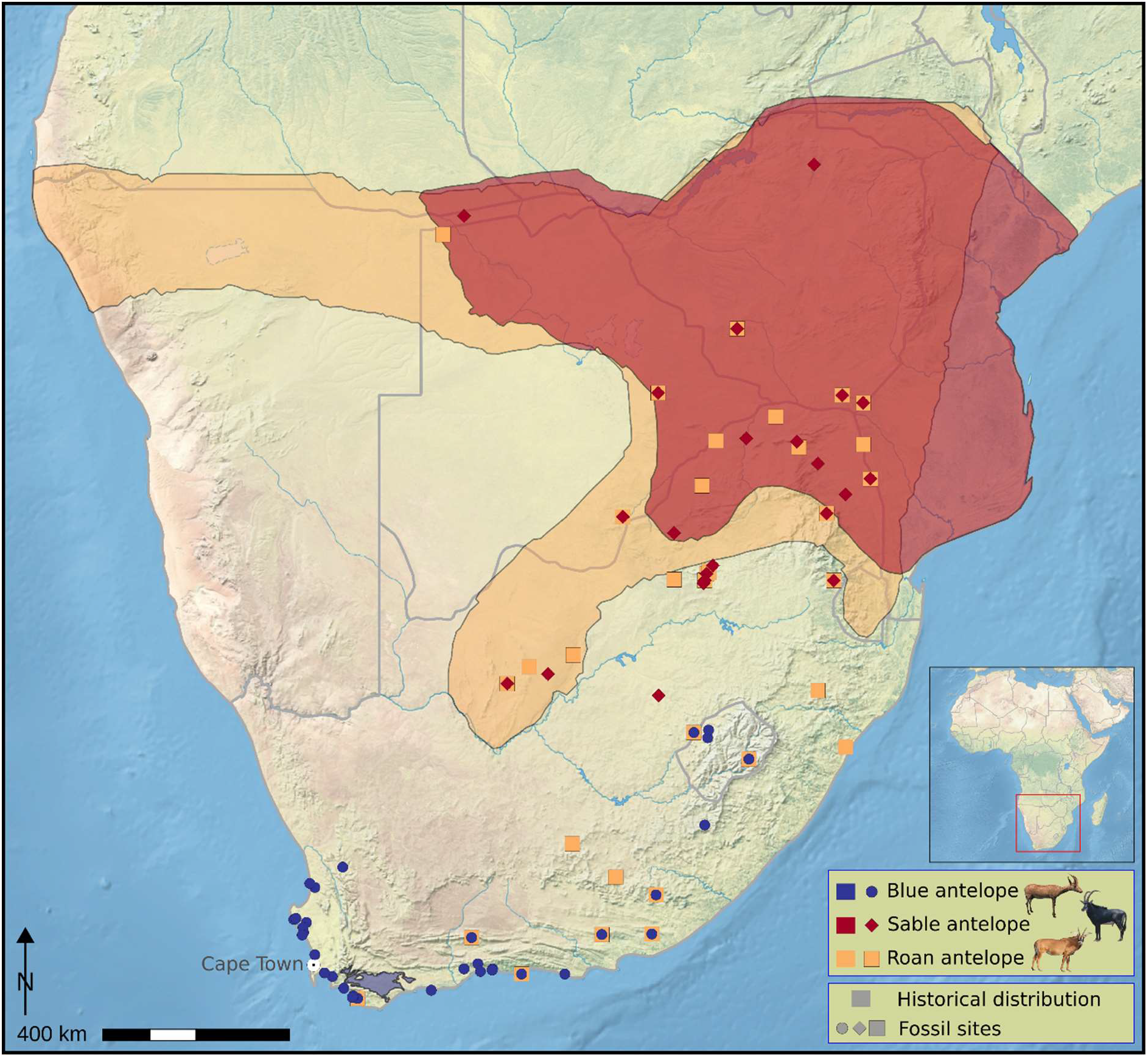
Species distribution and fossil sites. Historical distribution and Holocene and Pleistocene fossil sites of blue, sable and roan antelopes in southern Africa. The distributions of the roan and sable antelope further north are not shown (base map: https://www.naturalearthdata.com, prepared in QGIS v2.18 (https://qgis.org) (du Plessis 1969; Kerley et al. 2009; modified from Avery 2019). Photo credits: see fig. 2.

The genomic evidence for low diversity and therefore potentially low population sizes since the early Holocene confirms observations derived from the fossil record. Fossil evidence from southern Africa suggests that the blue antelope was both widespread and abundant during the last glacial period (~115 to 11.7 ka) (Klein 1987; Faith 2011). It became increasingly rare in the fossil record following the Pleistocene-Holocene transition (Faith 2011), with the long stratified sequence at Nelson Bay Cave – from where our fossil specimen originates – demonstrating that it remained a rare component of the large mammal community throughout the Holocene (Klein 1983). The onset of the population decline of the blue antelope near the Pleistocene-Holocene transition broadly coincides with the extinction of several other large-bodied grazers in southern Africa, including Cape zebra (*Equus capensis*), long-horn buffalo (*Syncerus antiquus*), and giant wildebeest (*Megalotragus priscus*), among others. These losses have been linked at least in part to declines in the availability and year-round productivity of grassy forage (Klein 1980; Brink and Lee-Thorp 1992; Faith 2014). This is particularly evident in the Cape Floristic Region, where grassy habitats diminished in conjunction with a rise in sea levels and drowning of the Palaeo-Agulhas Plain that would have contributed to dramatic habitat loss, disruption of possible migratory routes, and fragmentation of species ranges (Dingle and Rogers 1972; Klein 1983; van Andel 1989; Fisher et al. 2010; Compton 2011; Faith and Behrensmeyer 2013; Faith and Thompson 2013; Copeland et al. 2016; Venter et al. 2020). Rare Holocene occurrences of the blue antelope in the southwestern Cape (Klein and Cruz-Uribe 2016), the southern Cape (Klein 1983), and in the interior (Opperman 1987; Plug 1997), likely represent disjunct populations due to lack of dispersal corridors around the Cape Fold Mountains when sea levels were high (Compton 2011; Faith and Behrensmeyer 2013; Faith and Thompson 2013).

It appears that by the onset of historical times, the blue antelope had become restricted to the southern Cape (Klein 1983, but see Loubser et al. 1990), an area that provides limited foraging opportunities for large-bodied grazers (Klein 1983; Kerley et al. 2009; Faith and Behrensmeyer 2013; Cawthra et al. 2015; Cawthra et al. 2020). Thus, the environmental changes that marked the transition from the Pleistocene to the Holocene are likely to have resulted in a geographically isolated and small population (Skead 1980; Kerley et al. 2009), which conforms with our finding of very low nuclear diversity compared to the two extant *Hippotragus* species that were and are more broadly distributed across southern Africa (figs. 5 and 6). Archaeological evidence suggests that prehistoric human populations increased in southern Africa beginning ~15,000 years ago (Deacon and Thackeray 1984; Wadley 1993), but whether this contributed to the low population size of blue antelope by the early Holocene (e.g., via increased predation pressure) is presently uncertain. Linking prehistoric human activities to the blue antelope and other large-bodied ungulates from the region will require a more detailed understanding of their population histories through time and in relation to palaeoclimatic, palaeoenvironmental, and archaeological records.

Low nuclear diversity, and likely low population size, have characterised the blue antelope at least since the early Holocene, raising the question of whether the blue antelope was doomed to extinction, or whether it could have survived in the absence of increased human impacts in historical times. As Kerley et al. (2009) pointed out, due to increased susceptibility to drift there might have been an accumulation of mildly deleterious mutations in the small remaining blue antelope population, which might have put the species on a slow path to extinction.

Our finding of low diversity since the early Holocene, suggests that low diversity was most likely the result of climatically induced habitat fragmentation at the Pleistocene-Holocene transition, long before the introduction of livestock to the southern Cape (~2,000 years ago, Coutu et al. 2021) or overhunting and landscape transformation during the colonial era. Generally, low diversity is associated with increased extinction risk (DeWoody et al. 2021; Kardos et al. 2021). However, the persistence of blue antelope for at least 10,000 years despite the possibility of continuously low diversity suggests that low diversity might not have impacted the survivability of the species, similar to other species in southern Africa (Westbury et al. 2018; Sánchez-Barreiro et al. 2021). Cycles of habitat restrictions through past changes in sea level and forage availability may have led to repeated declines in population size and genetic diversity. Continuously low population sizes may have enabled more efficient purging of highly deleterious mutations by exposing them more often in a homozygous state, meaning that low population size might be an adaptive advantage rather than detriment (Xue et al. 2015; Dussex et al. 2021; Liu et al. 2021). However, hunting with firearms and land transformation in the blue antelope’s restricted habitat during the colonial era (FitzSimons 1920; Harper 1945; Kerley et al. 2003; Skead 2011; Faith and Thompson 2013) might have proved too much given this species’ low population size, ultimately culminating in its extinction at the hands of humans.

In conclusion, our study shows that using genomic data from fossil specimens from Africa opens up new research possibilities to understand evolutionary dynamics. For the blue antelope, we found low diversity throughout the Holocene, confirming observations derived from the fossil record. In the future, it will be interesting to see if genomic data from Pleistocene specimens will continue to reflect the patterns seen in the fossil record. Our results suggest that humans in the colonial era caused the extinction of a species that was likely already vulnerable due to habitat loss and range fragmentation at least since the onset of the Holocene.

## Materials and Methods

### Samples

We obtained tooth root or bone fossil samples from 25 blue antelope (*Hippotragus leucophaeus*) specimens from Archaeology Unit, Iziko Museums of South Africa, Cape Town, South Africa, ranging between 5,000–71,000 years BP (supplementary table S1) originating from six different fossil sites: Boomplaas Cave, Byneskranskop 1, Die Kelders Cave 1, Elands Bay Cave, Nelson Bay Cave, Klasies River Mouth 1/1A (fig. 1). Two samples were undated. Sample BNK1 O25/2 5.2 was radiocarbon-dated at the ^14^Chrono Centre, Queens University Belfast, Ireland using CALIB REV 7.0.0 (Stuiver and Reimer 1993, supplementary table S1). In addition, we used the single-stranded library of a historical sample from the Swedish Museum of Natural History in Stockholm, Sweden, from a previous study (Hempel et al. 2021a).

The Nelson Bay Cave specimen (NBC RB4 D3) that yielded a palaeogenome is a lower left deciduous premolar (dP_4_). The strongly pinched lingual lobes and presence of basal pillars are consistent with *Hippotragus*, and its relatively small size relative to a handful of larger *Hippotragus* dP_4_s from the site support attribution to *H. leucophaeus* (see Klein 1974; Faith 2012; supplementary fig. S7, supplementary table S18). It is derived from stratum RB, which is associated with AMS radiocarbon dates of 8,447 ± 39 ^14^C yr BP (9,545–9,460 cal yr BP) and 8,550 ± 39 ^14^C yr BP (9,520–9,305 cal yr BP) (Loftus et al. 2016). A Bayesian age model for the stratigraphic sequence at Nelson Bay Cave suggests the material derived from stratum RB broadly dates to between 9,800 cal yr BP and 9,300 cal yr BP (Loftus et al. 2016). The associated artefacts belong to the Later Stone Age Oakhurst industry, which is a flake-based technology with few microliths or formal lithic tools (Deacon 1984). In contrast to the Pleistocene levels at Nelson Bay Cave, blue antelope are relatively uncommon in RB, with the dominant ungulates including Cape grysbok (*Raphicerus melanotis*), bushbuck (*Tragelaphus scriptus*), and bushpig (*Potamochoerus larvatus*) (Klein 1972, 1983), all of which presumably favour more closed habitats than expected for blue antelope.

### Laboratory procedures

#### DNA preparation

We extracted DNA from all fossil samples following Dabney et al. (2013), using columns of Roche’s High Pure Viral Nucleic Acid Kit for purification. Subsequently, we built single-stranded libraries from these extracts according to Gansauge et al. (2017), employing an additional initial 15 min. incubation step with 0.5 μl USER Enzyme for Uracil removal (modified from Meyer et al. 2012). We used a maximum of 13 ng DNA as input for library construction. To determine the optimal number of amplification cycles for the subsequent dual-indexing PCR, we performed a qPCR (Thermo Scientific PikoReal Real-Time PCR System). Extraction and library blanks were run alongside all samples to check for the presence of contamination. All pre-PCR lab work was conducted in dedicated ancient DNA (aDNA) facilities at the University of Potsdam, Germany. The single-stranded library of the historical sample NRM 590107 was prepared in the same way as described above (Hempel et al. 2021a). All libraries from fossil specimens were test-sequenced on an Illumina NextSeq500 using custom primers (Gansauge and Meyer 2013; Paijmans et al. 2017) at the University of Potsdam to determine their endogenous DNA content. Only one sample from Nelson Bay Cave (NBC RB4 D3, dP_4_, fig. 1C) yielded sufficient endogenous DNA content for deeper sequencing. Therefore, we sequenced the library of NBC RB4 D3 using custom primers in one run on an Illumina NextSeq500 at the University of Potsdam, generating 75 bp single-end reads, and in another run on a NovaSeq6000 at the SciLifeLab, Sweden, generating 100 bp paired-end reads. We further sequenced the library of NRM 590107 using custom primers in three runs on an Illumina NextSeq500 at the University of Potsdam, generating 75 bp singleend reads, and on one run on a NovaSeq6000 at the SciLifeLab, generating 100 bp paired-end reads. In addition, for NRM 590107, we used 75 bp single-end read data generated from one run in Hempel et al. (2021a) (SRRxxxxxxx, supplementary table S2).

### Bioinformatic procedures and analyses

#### Nuclear genome

##### Data preprocessing and read mapping

We processed single- and paired-end reads separately prior to duplicate removal but combined all single-end runs of NRM 590107 before processing. We used Cutadapt v2.8 (Martin 2011) to trim Illumina adapter sequences (1 bp overlap) and to remove reads shorter than 30 bp. We merged paired-end reads with FLASH v1.2.11 (Magoč and Salzberg 2011) using a maximum overlap of 100 bp and discarded all unmerged reads. Subsequently, we mapped the resulting reads to the nuclear genome of the scimitar-horned oryx (*Oryx dammah*) (https://www.dnazoo.org/assemblies/Oryx_dammah, Humble et al. 2020) using the BWA aln algorithm v0.7.17 (Li and Durbin 2009) and default settings. For quality filtering, we used SAMtools view v1.10 (Li et al. 2009) to remove reads with a mapping quality of <30, discarding unmapped reads, and then sorted bam files with SAMtools sort. We then merged all reads from a single individual with SAMtools merge and performed duplicate removal with Picard MarkDuplicates v2.22.4 (Picard Toolkit 2020 - http://broadinstitute.github.io/picard). We ran mapDamage v2.2.0 (Jónsson et al. 2013) and rescaled the bam file using the rescale option (--rescale) to decrease the quality of misincorporations that are likely caused by aDNA damage.

We used published raw sequencing read data of three roan (*Hippotragus equinus*) (Gonçalves et al. 2021), eight sable antelopes (*Hippotragus niger*) (Koepfli et al. 2019) from contemporary samples, and the scimitar-horned oryx (*Oryx dammah*) (Humble et al. 2020) (supplementary table S4) in our analyses. The reads for these samples were treated in the same way as described above with the exception that merged and unmerged reads were both mapped, that the maximum overlap parameter in FLASH v1.2.11 was adjusted according to sequencing cycle length and that no rescaling was performed. For one roan antelope sample (10954) and the scimitar-horned oryx sample (SB20612), both from 10X Genomics libraries, 22 bp were trimmed from R1 after adapter trimming with Cutadapt v2.8 using FASTA/Q Trimmer from the FASTX-Toolkit v0.0.14 (http://hannonlab.cshl.edu/fastx_toolkit).

Since sex chromosomes and mitochondrial genomes have different inheritance patterns relative to autosomes and could bias downstream analyses, we removed 53 scaffolds that were identified to likely represent X and Y chromosomes and the mitochondrial genome in the scimitar-horned oryx reference genome (Humble et al. 2020; Hempel et al. 2021b).

As input fasta files for *D* statistics (Green et al. 2010; Durand et al. 2011), the nuclear diversity estimates and Bayesian molecular dating, we generated pseudohaploid consensus sequences for all specimens using Consensify (Barlow et al. 2020, https://github.com/jlapaijmans/Consensify). We generated the base count input file using ANGSD v0.923 (Korneliussen et al. 2014) (-minQ 30, -minMapQ 30, -uniqueonly 1, -remove_bads 1, -baq 2, -dumpCounts 3, -C 0, -only_proper_pairs 0, -doCounts 1, -trim 0). For the fossil and historical blue antelope specimens, the rescaled bam files from mapDamage were used as input files. We ran Consensify with twice the mean coverage of each sample as the maxDepth parameter and using only autosomal scaffolds larger than 1 Mb. To check if sequence coverage had an effect on downstream results, we generated an additional set of subsampled files for all individuals (fossil, historical and contemporary) with SAMtools view, setting the baseline to the sample with the lowest mean coverage (2.14x, NBC RB4 D3) before running Consensify.

For the sliding window tree analyses we generated pseudohaploid consensus sequences using random read sampling (-doFasta 1) in ANGSD and the following parameter settings: -minQ 30, -minMapQ 30, -uniqueonly 1, -only_proper_pairs 0, -remove_bads 1, -baq 2, -C 0, -explode 1, -doCounts 1, -trim 0, -basesPerLine 70. We only included autosomal scaffolds larger than 1 Mb (supplementary table S6). For the fossil and historical specimens of blue antelope, the rescaled bam files from mapDamage were used as input files.

###### Species tree inference

We constructed a neighbour-joining tree using the nuclear genome data from the two blue, three roan (Gonçalves et al. 2021) and eight sable antelopes (Koepfli et al. 2019), along with the scimitar-horned oryx (SB20612, Humble et al. 2020) as outgroup to examine the dominant phylogenetic signal. The rescaled bam files from mapDamage for the fossil and historical blue antelope specimens as well as the bam files for all contemporary individuals were used as input files. We generated the input distance matrix in ANGSD using a consensus base call approach (-doIBS 2), removing singletons (-minMinor 2) and transitions (-rmTrans 1), and using only positions where all individuals had coverage (-minInd 14) (-doCounts 1, -makeMatrix 1, -minQ 30, -minMapQ 30, -mininddepth 2, -uniqueonly 1, -only_proper_pairs 0, -remove_bads 1, -baq 2, -trim 0, -C 0, -doMajorMinor 1, -GL 2). We used FastME v2.1.6.1 (Lefort et al. 2015) to generate a neighbour-joining tree from the distance matrix using default parameters. The tree was visualised with FigTree v1.4.3 (https://github.com/rambaut/figtree).

To also infer the species tree while accounting for incomplete lineage sorting and gene flow, we ran a sliding window tree analysis using the tool WindowTrees v1.0.0 (https://github.com/achimklittich/WindowTrees) with fasta files generated from random read sampling as input to generate non-overlapping sliding windows as input to infer phylogenies per window using IQ-TREE v2.2.0 (Minh et al. 2020). We used binary mode to exclude transitions (--binary) and a missing data threshold of 50% (-N 0.5) with a window size of 20 kb (-w) and a gap size of 80 kb (-lw 80000) between windows. For this analysis only one individual of each species was used (roan: 10954, sable: SB2152, blue antelope: NBC RB4 D3). The scimitar-horned oryx was used as outgroup (SB20612) (--outgroup). We used the resulting phylogenies while adding the mitochondrial genomes (without control regions) as an additional window as input for ASTRAL v4.10 (Rabiee et al. 2019, https://github.com/smirarab/ASTRAL) to infer the species tree under the multi-species coalescent with local posterior probability branch supports. Gene- and site-concordance factor branch supports were also inferred using IQ-TREE2.

###### Fossil calibrated phylogeny

We fossil-calibrated the species phylogeny generated from the combined nuclear and mitochondrial data set. To avoid the impact of incomplete lineage sorting on molecular dating (Mendes and Hahn 2016), windows were filtered to include only those that had a concordant phylogenetic signal with the species tree, as inferred from the ASTRAL and neighbour-joining analyses (see above). Similarly, we avoided further phylogenetic biases by using trees with strong branch supports (bootstrap support >90) and low rate variation among lineages (CoV in root-to-tip length <0.1; Vankan et al. 2022). We used the following individuals: the early Holocene blue antelope (NBC RB4 D3), the roan antelope (10954, Gonçalves et al. 2021), the sable antelope (SB2152, Koepfli et al. 2019) and the scimitar-horned oryx (SB20612, Humble et al. 2020) as outgroup. As input files we used the fasta files generated with Consensify (see above) but including only transversions. For calibration we used a uniform prior between 3.6 and 4.5 Mya for the split between *Oryx* and *Hippotragus* (Vrba and Gatesy 1994; Deino 2011; Gentry 2011; for further explanations see Bibi 2013), with a soft maximum bound. We used approximate likelihood computation as implemented in MCMCtree (in PAML v4.9; dos Reis and Yang 2011) with an MCMC chain of 10M steps, discarding the first million as burn-in. We verified that the analysis reached convergence by verifying all parameter traces and replicating the analysis to confirm convergence to an identical optimum (supplementary files S3-10). The tree was visualised with FigTree.

###### *D* statistics

For *D* statistics analysis, we used the fasta files generated with Consensify of the roan and sable antelope individuals with the highest coverage (10954, Gonçalves et al. 2021, and SB2152, Koepfli et al. 2019) together with the scimitar-horned oryx as outgroup (SB20612, Humble et al. 2020) and either the fossil or the historical blue antelope specimen. We used the topology resulting from the 20 kb sliding window multi-species coalescent and neighbourjoining analyses (see above, fig. 2 and supplementary fig. S3) placing the sable antelope in position one, the blue antelope in position two, and the roan antelope in position three (fig. 3B). We conducted *D* statistics/abbababa test with the tool Dstat (transversion only version, https://github.com/jacahill/Admixture) and calculated the standard error using the weighted_block_jackknife_D tool with a 1 Mb window size. From this we calculated Z-scores with a Z-score>|3| defined as significant.

###### Inferences of gene flow directionality and timing

To investigate the direction and timing of gene flow we used a sliding window tree approach. As input we used the randomly sampled fasta files (see above). We then used the tool WindowTrees (https://github.com/achimklittich/WindowTrees) to generate non-overlapping sliding window maximum-likelihood phylogenies using RAxML v8.2.12 (Stamatakis 2014) in binary mode to exclude transitions (--binary), a window size of 100 kb (-w) and a threshold for missing data of 50% (-N 0.5). Prior to the introduction of the WindowTrees tool, this procedure was already successfully used to determine gene flow direction in other studies (Barlow et al. 2018; Westbury et al. 2020; Paijmans et al. 2021). This was performed for all possible combinations with two roan, one sable and one blue antelope using the following individuals: roan: 10954, He108: sable: SB2152, HN216; blue antelope: NBC RB4 D3, NRM 590107. The scimitar-horned oryx (SB20612) was defined as outgroup in each case (--outgroup).

In addition, we used the resulting phylogenies from the sliding window tree analysis that used WindowTrees in binary mode with a missing data threshold of 50%, a window size of 20 kb and a gap size of 80 kb (-lw 80000) between windows (see above). We examined the hypothesis of gene flow in the direction of blue into roan antelope using estimates of gene trees and their phylogenetic branch lengths. Under a scenario of gene flow from blue into roan antelope, the portion of introgressed loci will show a signal of more recent common ancestry between the roan and the sable antelope relative to the species tree topology. On the other hand, a scenario of gene flow in the direction from roan into blue antelope will lead all of the loci to have a signal of similar times of common ancestry between roan and sable antelope in accordance with the species tree topology (fig. 4A). To examine these hypotheses, we extracted the sum of phylogenetic branch lengths separating roan and sable antelope instead of roan and blue antelope as to avoid biases that may influence the branch length, e.g., possible DNA damage and higher sequencing errors, due to lower coverage, as expected from the fossil specimen. We then compared this signal between the gene trees in which roan and blue antelope are sister species (potentially including introgressed regions) against gene trees in which sable and blue antelope are sister species (showing distances between common ancestors of those species). To exclude extremely high and extremely low distances, which likely arise due to the high variance in estimates of branch lengths with negligible information available, we filtered out distances of <0.0002 and >0.9. For any occurrence of gene flow from the blue into the roan antelope, a bimodal distribution would have been expected due to the relatively shorter interspecific branch lengths that would arise. Genetic distances from gene trees were extracted using the R package *ape* v5.5 (Paradis and Schliep 2019, supplementary file S11).

###### Species-wide nuclear diversity comparisons

We performed within species pairwise comparisons to estimate the species-wide nuclear diversity for each species within the genus *Hippotragus*. We implemented an approach that we show can be used for low coverage data and take the patterns of unusual DNA damage found in blue antelope specimen NRM 590107 into account.

We used the fasta files generated with Consensify from the two blue, three roan, and eight sable antelope samples, and the single outgroup scimitar-horned oryx, and filtered those by excluding positions with missing data and positions where all individuals had the same allele (uninformative sites) using a custom perl script. We then performed pairwise difference estimates for the individuals of each species by counting the number of different alleles while excluding all potential aDNA damage (transitions) and all differences resembling the damage pattern found in NRM 590107 (guanine to thymine and cytosine to adenine and vice versa). The values were normalised by dividing them by the total number of positions after filtering. We plotted the values for each pairwise comparison together with the mean for roan and sable antelope. We additionally ran the analysis with the input files that were subsampled to 2.14x mean coverage before running Consensify to test for the effects of sequence coverage.

To examine how the number of differences between individuals compares to a more standard measure of diversity, i.e. heterozygosity, we calculated allele frequencies using genotype likelihoods for each roan and sable antelope individual using ANGSD, setting the -setMaxDepth parameter to twice the mean coverage and with the following parameter settings: -setMinDepthInd 5, -minInd 1, -doCounts 1, -GL 2, -doSaf 1, -fold 1, -minQ 30, -minMapQ 30, -uniqueonly 1, -remove_bads 1, -only_proper_pairs 0, -trim 0, -C 0, -baq 2. Next, we calculated the site frequency spectrum using realSFS in ANGSD. This analysis was performed once using the full dataset and once with the bam files subsampled with SAMtools view to the lowest mean coverage of a specimen in the roan/sable antelope data set (4.26x of sable antelope SB1954) to ensure no bias could result from uneven coverage. We plotted the values for the subsampled calculations per individual together with the mean for each species.

##### Mitochondrial genome

###### Read mapping

We treated the reads for the mitochondrial genome of NBC RB4 D3 in the same way as described for the nuclear genome but used the mitochondrial genome of the blue antelope (MW222233, Hempel et al. 2021a) as reference and removed duplicates with MarkDupsByStartEnd (https://github.com/dariober/Java-cafe/tree/master/MarkDupsByStartEnd). We generated a consensus sequence in Geneious R10 v10.2.3 (Kearse et al. 2012, https://www.geneious.com) using an 85% majority rule threshold for base calling, a minimum coverage of 3x and the “trim to reference” option. To improve the coverage at the end of the sequence, we mapped and filtered the data again as described above, but used a reference with 200 bp shifted from the end to the start of the reference, considering the circular nature of the mitochondrial genome. We called a consensus sequence as described before and subsequently moved the 200 bp back to the end of the sequence. Then we called a consensus sequence from both sequences using a 50% majority rule threshold for base calling (option “50%—Strict: Bases matching at least 50% of the sequences”).

We generated mitochondrial genomes from the raw sequencing data of all roan and sable antelope specimens and the outgroup scimitar-horned oryx specimen for which assembled mitogenomes were not available (roan: He95, He108, 10954; sable antelope: HN216, HN250; scimitar-horned oryx: 20612) (supplementary table S5). We treated the data in the same way as described for the nuclear genomes while using conspecific mitochondrial genomes as references (roan and sable antelope and scimitar-horned oryx: JN632647, JN632648, JN632677; Hassanin et al. 2012) and employing a second mapping step as described for NBC RB4 D3 but varying the length of the reference part that was moved for the second mapping (see above) to be twice the sequencing cycle length of each sample.

###### Maximum-likelihood phylogenies from mitochondrial genomes

We built an alignment with the consensus sequences of the two blue, three roan and eight sable antelopes using the MAFFT algorithm v7.450 (Katoh et al. 2002; Katoh and Standley 2013) and default parameters as implemented in Geneious. We used the scimitar-horned oryx as outgroup (SB20612). We removed the control region from all sequences prior to aligning them due to their limited alignability for different species (supplementary files S12 and S13). We generated two maximum-likelihood phylogenies with 1000 bootstrap replicates each using RAxML v8.2.12, once with the GTR+G substitution model and the other with the BINGAMMA model (Stamatakis 2014). For the latter, the alignment was first transformed into binary format to only score transversions. The tree was visualised with FigTree.

###### Haplotype network with mitochondrial genomes

To determine the extent of differences between our fossil and the previously published historical specimens of the blue antelope, we aligned the mitochondrial genome of NBC RB4 D3 to the other two available complete mitochondrial blue antelope genomes (MW222233, MW222234, Hempel et al. 2021a, supplementary table S17) using the MAFFT algorithm with default settings as implemented in Geneious. We removed all ambiguities/missing data (supplementary files S14 and S15). Subsequently, we constructed a median-joining network with POPART v1.7 (Bandelt et al. 1999; Leigh and Bryant 2015) and calculated the number of segregating sites and nucleotide diversity. In addition, we aligned the mitochondrial genome of NBC RB4 D3 with the two available complete and the two partial mitochondrial genomes of the blue antelope (MW228401, MW228402, Hempel, et al. 2021a, supplementary table S17) using again the MAFFT algorithm with default settings as implemented in Geneious and repeated all steps described for the alignment with the complete mitochondrial genomes to generate a median-joining network and determine the number of segregating sites and the nucleotide diversity using POPART.

## Supplementary Material

Supplementary data are available at Molecular Biology and Evolution online including alignments for the haplotype networks and maximum-likelihood phylogenies and the R scripts for the sliding window tree genetic distance analysis and the phylogenomic analysis.

## Data availability

The BioProject number of this project in GenBank is PRJNAxxxxxx. The complete mitochondrial genome of the fossil blue antelope specimen NBC RB4 D3 is available at GenBank under the accession number xxxxxx. The untrimmed raw data were uploaded for the fossil specimen NBC RB4 D3 to the Sequence Read Archive under SRRxxxxx, SRRxxxxxx, SRRxxxxxx, SRRxxxxxx and SRRxxxxxxx and for the historical specimen NRM 590107 under SRRxxxxx–SRRxxxxx and SRRxxxxxx. For NRM 590107 run SRRxxxxxx from BioProject PRJNAxxxxxx was used as well.

## Acknowledgments

This work was supported by the Deutsche Forschungsgemeinschaft (DFG, German Research Foundation) (project number 315696891, HO 3492/3-1, BI 1879/2-1 to M.H. and F.B.) and by the Swedish Research Council (2017-04647) and FORMAS (2018-01640 both to L.D.). We would like to acknowledge the support of Iziko Museums of South Africa, Heritage Western Cape, Cape Nature, Eastern Cape Provincial Heritage Resources Authority and South African Heritage Resources Agency. We would like to thank Wendy Black, Wilhelmina Seconna and Mark De Benedictis for access to and support while at the Archaeology Unit, Iziko Museums of South Africa. We thank Margarida Gonçalves and Raquel Godinho for providing the raw data for the roan antelope sample 10954 and Gaik Tamazian and Pavel Dobrynin for providing the raw data for the sable antelope samples HN216 and HN250. We would like to thank Stefanie Hartmann for providing the custom perl script for data filtering and Michaela Preick for lab work in screening samples. The authors also acknowledge support from Science for Life Laboratory, the Knut and Alice Wallenberg Foundation, the National Genomics Infrastructure funded by the Swedish Research Council, and Uppsala Multidisciplinary Center for Advanced Computational Science for assistance with massively parallel sequencing and access to the UPPMAX computational infrastructure.

## Author Contributions

The study was designed by E.H., F.B., J.T.F., J.S.B., M.H. and M.V.W. Funding was acquired by F.B. and M.H. J.T.F. identified and measured specimens. E.H. sampled the specimens and performed lab work. E.H., D.A.D. and M.V.W. performed DNA analyses with input from M.H. A.M.K. wrote the WindowTrees program. E.H. and M.V.W. wrote the manuscript with input from F.B., J.T.F., K.-P.K., D.A.D., D.C.K. and M.H. Resources were supplied by L.D. and M.H. Final editing and manuscript preparation was coordinated by E.H. All contributing authors read and agreed to the final manuscript.

