## Supplementary file S1 for "Blue turns to grey - Palaeogenomic insights into the evolutionary history and extinction of the blue antelope (*Hippotragus leucophaeus*)"

#### Supplementary figures

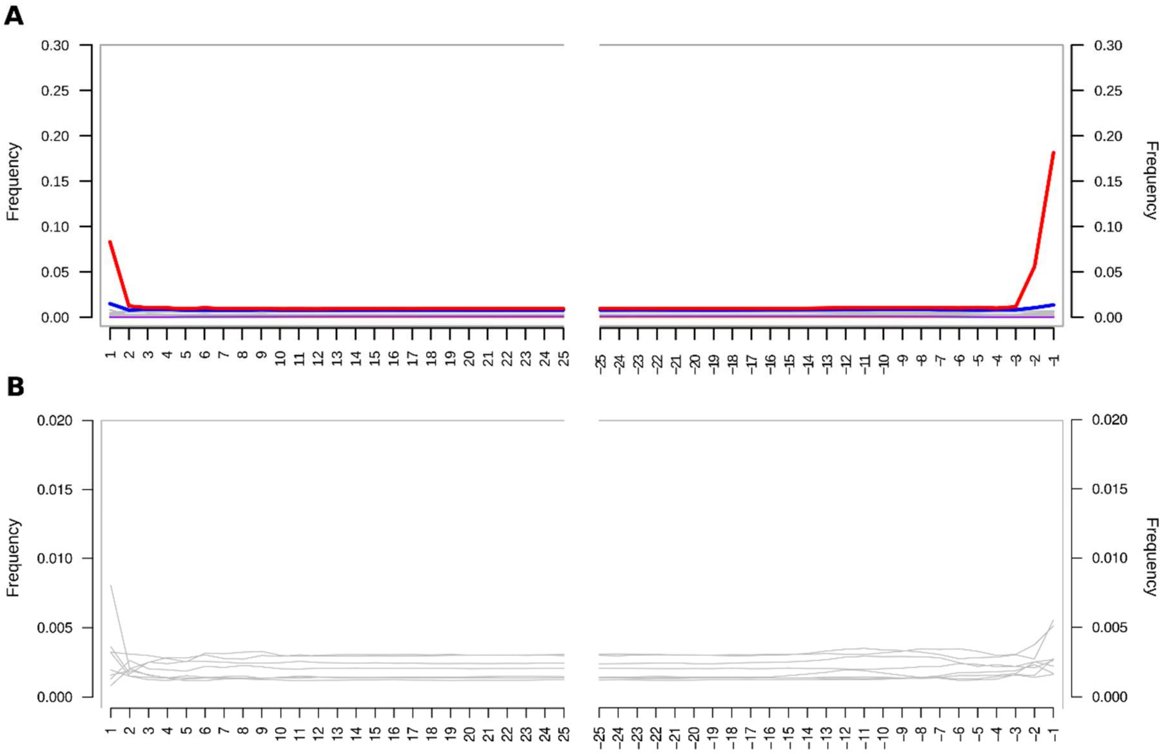

**Supplementary fig. S1.** DNA damage patterns for blue antelope specimen NBC RB4 D3 using mapDamage v2.2.0 (Jónsson et al. 2013). (**A**) MapDamage output, red: cytosine to thymine substitutions, blue: guanine to adenine substitutions, grey: all other substitutions. As expected from sequences from single-stranded library preparation, no guanine to adenine substitutions are shown on the 3' end (right) (Meyer et al. 2012) (**B**) Modified mapDamage output displaying only transversions. No transversion is distinctly higher. Note the different scales on the y-axes in A and B.

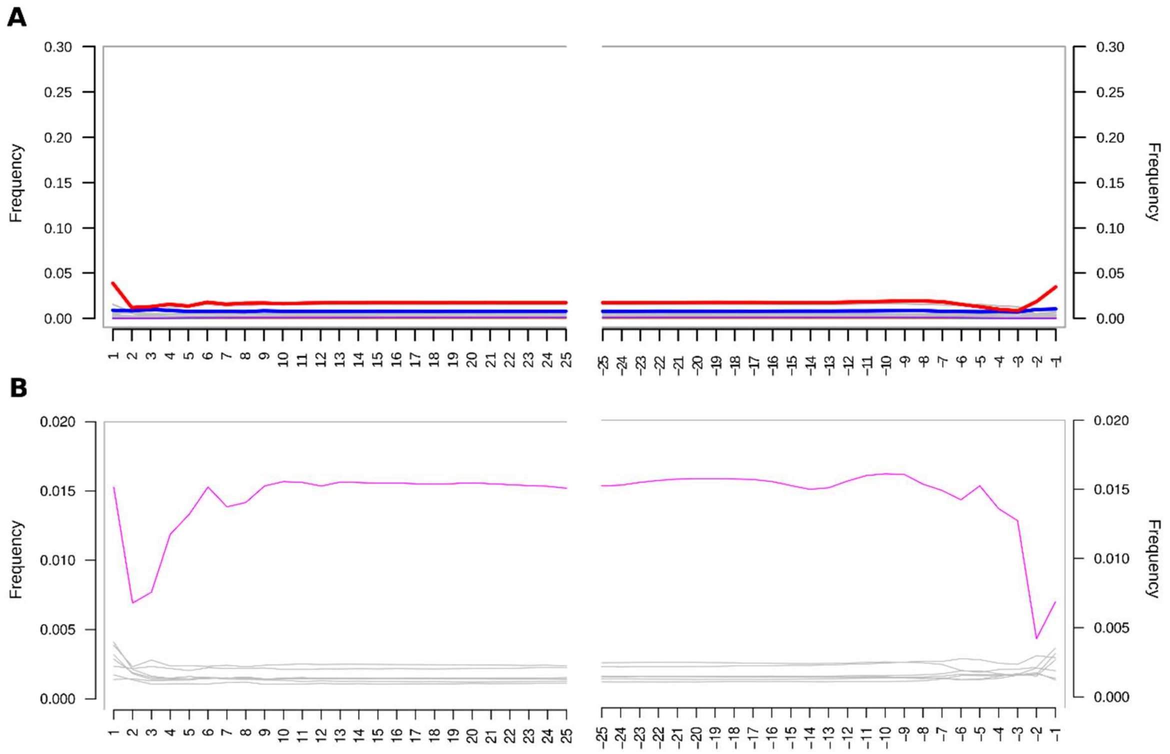

**Supplementary fig. S2.** DNA damage patterns for blue antelope specimen NRM 590107 using mapDamage v2.2.0 (Jónsson et al. 2013). (**A**) MapDamage output, red: cytosine to thymine substitutions, blue: guanine to adenine substitutions, grey: all other substitutions. As expected from sequences from single-stranded library preparation no guanine to adenine substitutions are shown on the 3' end (right) (Meyer et al. 2012) (**B**) Modified mapDamage output displaying only transversions, showing elevated guanine to thymine levels (pink). Patterns like this could originate from hydrogen peroxide treatment of the sample (Kvam and Tyrrell 1997; Nohmi et al. 2005). Note the different scales on the y-axes in A and B.

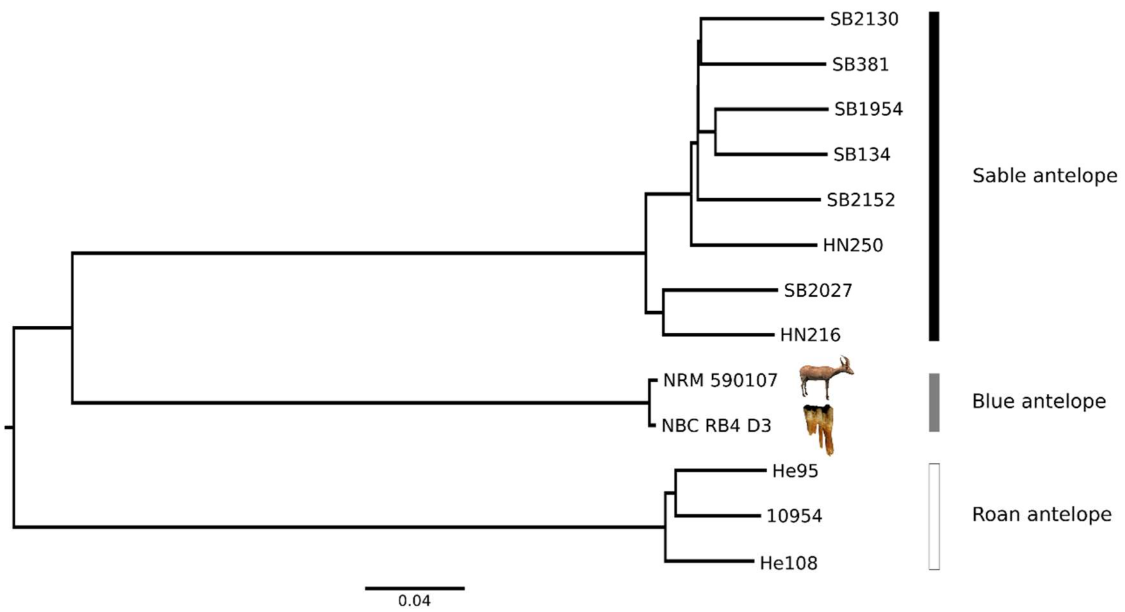

**Supplementary fig. S3.** Neighbour-joining phylogenetic tree (singletons removed) of the three *Hippotragus* species based on nuclear genomes showing the sister relationship of blue and sable antelopes. The phylogeny was constructed using ANGSD v0.923 (Korneliussen et al. 2014) and FastME v2.1.6.1 (Lefort et al. 2015). The scimitar-horned oryx (SB20612, Humble et al. 2020) was used as outgroup (not shown). Roan and sable antelope raw data are from Gonçalves et al. (2021) and Koepfli et al. (2019). Photo credits: NBC RB4 D3: J.T. Faith, courtesy: Archaeology Unit, Iziko Museums of South Africa; NRM 590107: Hempel et al. 2021a.

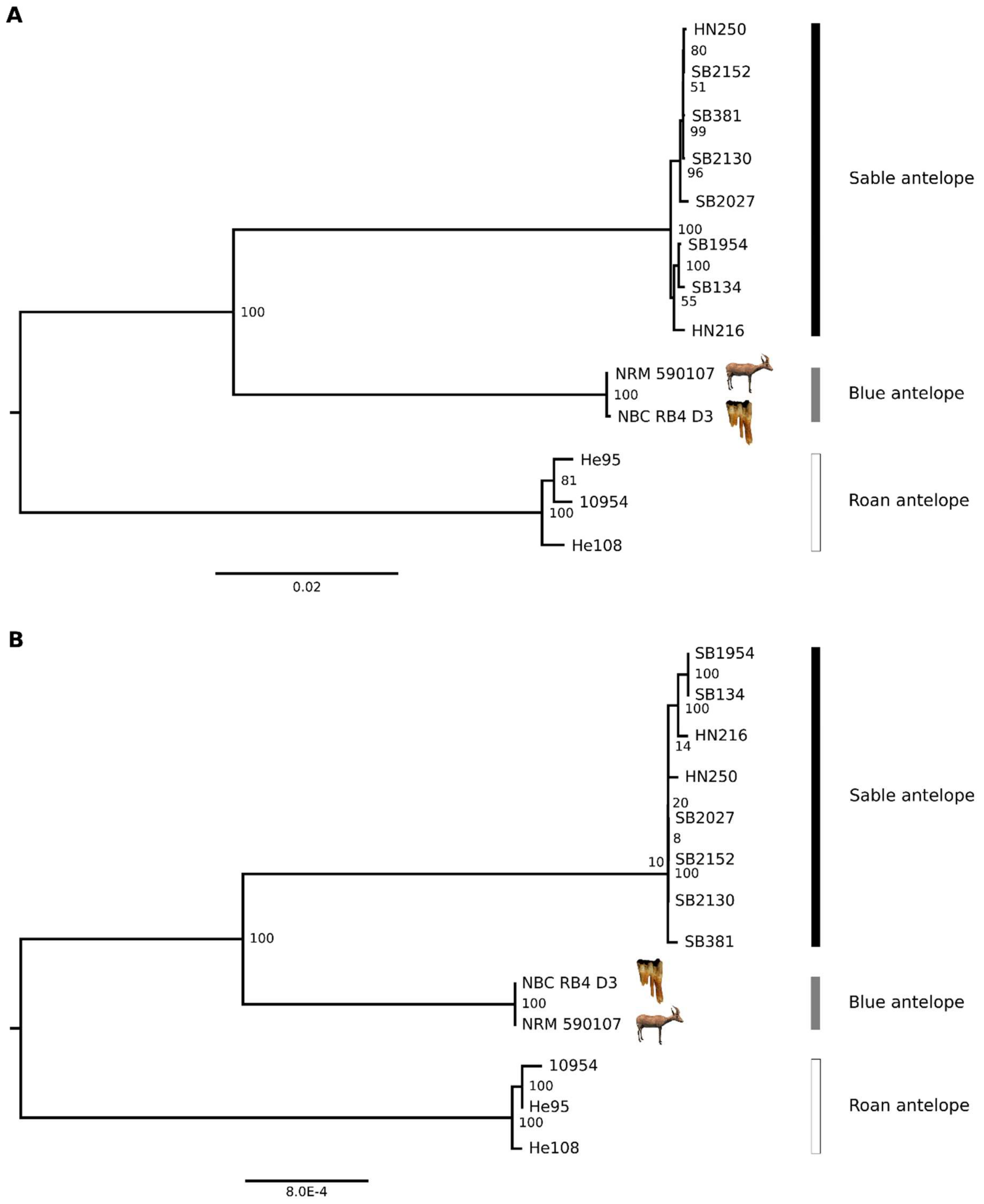

**Supplementary fig. S4.** Mitochondrial phylogenetic analyses. (**A**) Maximum-likelihood phylogeny of mitochondrial genomes of the three *Hippotragus* species. The phylogeny was generated using RAxML v8.2.12 with a GTR+G substitution model (Stamatakis 2014). The control region was excluded from the alignment (15,440 bp alignment length). (**B**) Maximum-likelihood phylogeny of mitochondrial genomes of the three *Hippotragus* species. The phylogeny is based on transversions only and was generated using RAxML with a BINGAMMA substitution model. The control region was excluded from the alignment (15,440 bp alignment length, non-binary format). Node labels show bootstrap support from 1000 replicates. In both analyses, the scimitar-horned oryx (*Oryx dammah*, SB20612, Humble et al. 2020) was used as the outgroup (not shown). Mitochondrial genome sequences of roan and sable antelope generated from Gonçalves et al. (2021) and Koepfli et al. (2019). Photo credits: see supplementary fig. S3.

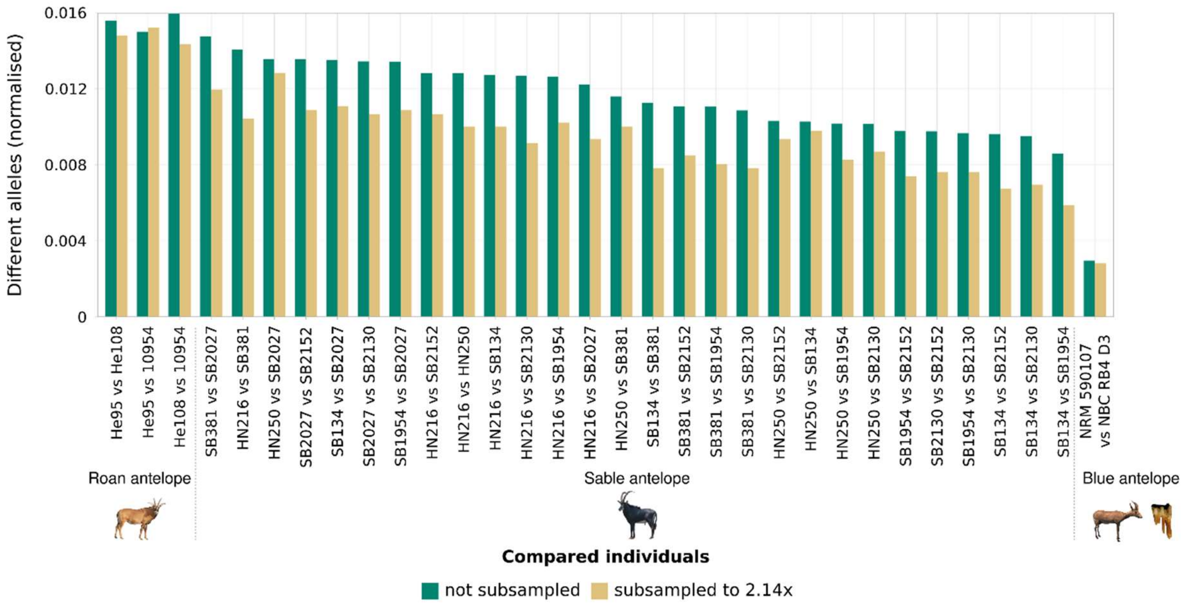

**Supplementary fig. S5.** Species-wide nuclear genome diversity of the three *Hippotragus* species computed from pairwise comparisons for data subsampled to 2.14x mean coverage (yellow bars) and data not subsampled (green bars). Data normalised for 4,596 and 2,243,953 sites, respectively. Genome data of roan and sable antelope are from Gonçalves et al. (2021) and Koepfli et al. (2019), respectively. Photo credits: NBC RB4 D3: J.T. Faith, courtesy: Archaeology Unit, Iziko Museums of South Africa; NRM 590107: Hempel et al. 2021a, roan antelope: Charles J. Sharp, wikimedia commons, CC-BY 4.0; sable antelope: Paulmaz, wikimedia commons, CC-BY 3.0.

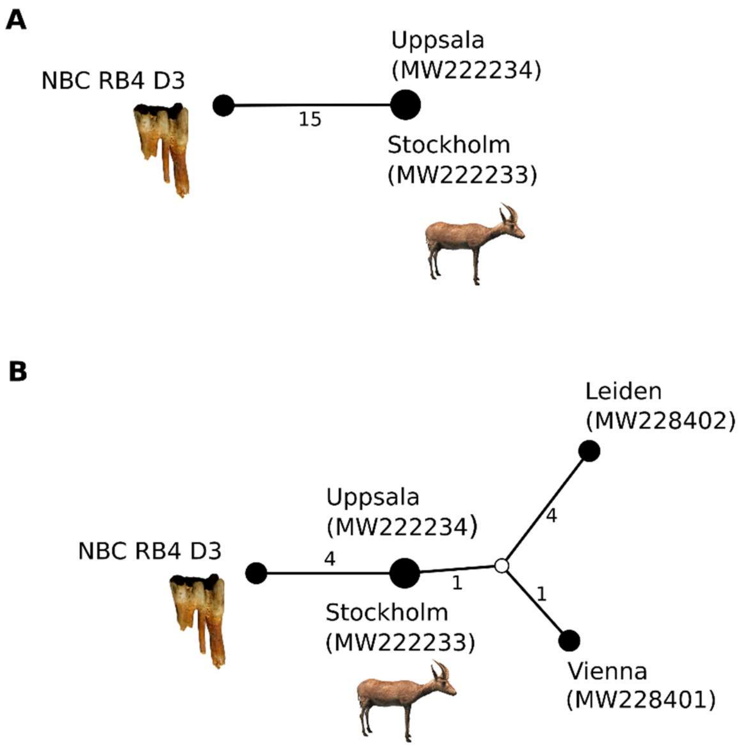

**Supplementary fig. S6.** Blue antelope mitochondrial haplotype networks. (**A**) Median-joining network of three complete blue antelope mitochondrial genomes (alignment length 16,492 bp) generated in POPART v1.7 (Bandelt et al. 1999; Leigh and Bryant 2015). Ambiguities/missing data were excluded. Numbers at lines represent mutational steps. Circle sizes correspond to frequency of haplotypes. City names are locations of the museum specimens, GenBank accession numbers are in parentheses. (**B**) Median-joining network of three complete and two partial mitochondrial genomes (alignment length 6,300 bp). The white circle represents an unsampled haplotype. Photo credits: see supplementary fig. S3.

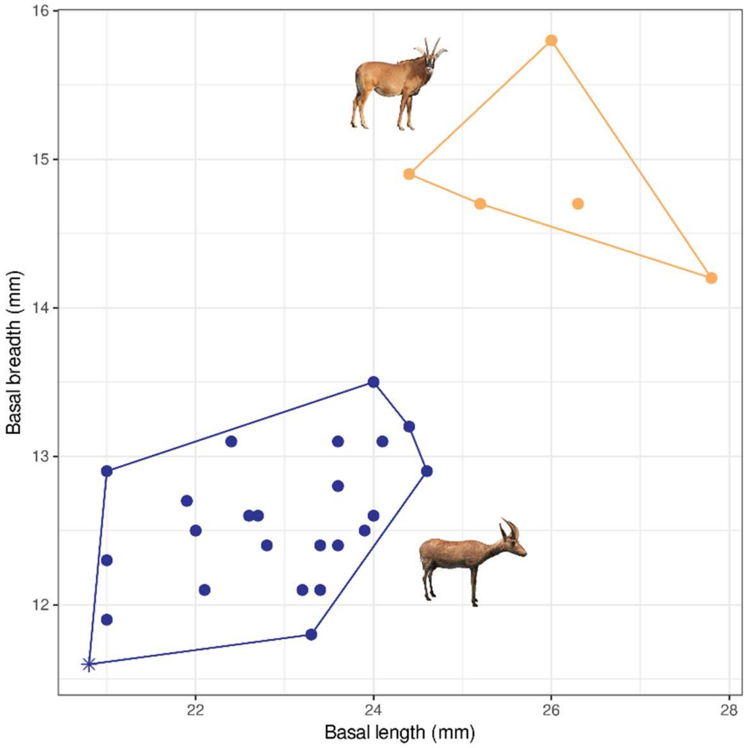

**Supplementary fig. S7.** Bivariate plot of basal length versus basal width of blue (indicated in dark blue) and roan antelope (indicated in orange) $dP_4$s from late Quaternary archaeological sites in the Western Cape, South Africa. Roan antelope specimens are from Boomplaas Cave and Nelson Bay Cave; blue antelope specimens are from Boomplaas Cave, Blombos Cave, Die Kelders Cave 1, Nelson Bay Cave, and Klasies River Mouth 1 (see supplementary table S18). Length and width measurements (in mm) taken at the base of the tooth crown. Nelson Bay Cave specimen NBC RB4 D3 is indicated with an asterisk. Photo credits: see supplementary fig. S5.

### Supplementary tables

**Supplementary table S1.** Sample information for the fossil blue antelope (*Hippotragus leucophaeus*) specimens housed at the Archaeology Unit, Iziko Museums of South Africa, Cape Town, South Africa, from which DNA was extracted in this study. The sample from which DNA was extracted successfully is printed in blue.

| Site | Sample stratification number | Species identification | Age of layer [years BP] | Sampled material | References |
|---|---|---|---|---|---|
| Elands Bay Cave | EBC KM E6 | *Hippotragus leucophaeus* | 12,000–14,000 | Tooth root ($P^4$) | Klein and Cruz-Uribe 2016; Parkington 2016 |
| Die Kelders Cave 1 | DK1 M/J D3 | *Hippotragus* cf. *leucophaeus* | 57,000–71,000 | Tooth root ($I_2$) | Klein and Cruz-Uribe 2000; Marean et al. 2000 |
| | DK1 MJA E5 | *Hippotragus* cf. *leucophaeus* | 57,000–71,000 | Tooth root ($I_1$) | |
| Byneskranskop 1 | BNK1 O29 7 | *Hippotragus leucophaeus* | 5,900–6,200 | Mandible fragment | Schweitzer and Wilson 1982; Loftus et al. 2016 |
| | BNK1 O29 5 | *Hippotragus leucophaeus* | 6,000–6,300 | Tooth root ($P^3$) | |
| | BNK1 P29 19 QUI | *Hippotragus leucophaeus* | 16,000–17,000 | Tooth root ($P_4$) | |
| | BNK1 O25/2 5.2 | *Hippotragus leucophaeus* | $^{14}C$ 3881 ± 42[1] | For aDNA extraction: tooth root ($M^2$) & for carbon dating: maxilla | |
| Boomplaas Cave | BPA BGM CL3 016/61 4.4890 | *Hippotragus leucophaeus* | 13,700–17,800 | Mandible | Klein 1978; Pargeter et al. 2018 |
| | BPA BRL7 Q15/51 3.5127 | *Hippotragus leucophaeus* | 12,600–14,000 | Mandible | |
| | BPA LPM P13 6.419 | *Hippotragus leucophaeus* | 21,400–23,500 | Mandible | |

| | | | | | |
|---|---|---|---|---|---|
| Nelson Bay Cave | NBC BSBI 9 D9 | *Hippotragus leucophaeus* | 6,200–7,000 | Mandible | Klein 1983; Loftus et al. 2016 |
| | NBC RA 9 B1 | *Hippotragus leucophaeus* | 6,900–9,200 | Mandible | |
| | NBC RB 4 D3 | *Hippotragus leucophaeus* | 9,300–9,800 | Tooth root ($dP_4$) | |
| | NBC J 1 D5 | *Hippotragus leucophaeus* | 9,500–11,400 | Tooth root ($M^3$) | |
| | NBC BSJ 80 B4 | *Hippotragus leucophaeus* | 10,900–11,900 | Mandible | |
| | NBC CS 2 D5 | *Hippotragus leucophaeus* | 11,700–12,200 | Mandible | |
| | NBC CS 52 D8 | *Hippotragus leucophaeus* | 11,700–12,200 | Mandible | |
| | NBC BSTL 26 D3 | *Hippotragus leucophaeus* | 12,000–14,900 | Tooth root ($M^1$) | |
| | NBC GBSL 43 | *Hippotragus leucophaeus* | 12,000–14,900 | Tooth root ($P^2$) | |
| | NBC YSL 8 G8 | *Hippotragus leucophaeus* | 16,300–22,100 | Tooth root ($M_1$) | |
| | NBC YGL 20 D5 | *Hippotragus leucophaeus* | 21,000–23,500 | Maxilla | |
| | NBC DS 18 1 | *Hippotragus leucophaeus* | Stratigraphic position unclear | Mandible | |
| Klasies River Mouth 1/1A | KRM1 13 ??? 2071 | *Hippotragus leucophaeus* | ~60,000–65,000 | Tooth root ($P^4$) | Klein 1976; Singer and Wymer 1982; Grine et al. 2017 |
| | KRM1A 9C 41459 | *Hippotragus leucophaeus* | ~60,000 | Mandible | |
| | KRM1A 6 1A 40.42 | *Hippotragus leucophaeus* | ~60,000 | Tooth root ($M_2$) | |

[1] Radiocarbon dated in this study calibrated using Reimer et al. (2013) and CALIB REV 7.0.0 (Stuiver and Reimer 1993)

**Supplementary table S2.** Paired-end (PE) and single-end (SE) sequencing results from the fossil (NBC RB4 D3) and historical (NRM 590107) blue antelope (*Hippotragus leucophaeus*) samples mapped to the nuclear genome assembly of the scimitar-horned oryx (*Oryx dammah*, https://www.dnazoo.org/assemblies/Oryx_dammah, Humble et al. 2020). Only merged reads were used. The paired-end run of NBC RB4 D3 was split by the Sequence Read Archive into four files.

see supplementary file S2

**Supplementary table S3.** Paired-end (PE) and single-end (SE) sequencing results from the fossil (NBC RB4 D3) blue antelope sample mapped to the mitochondrial genome of the blue antelope (*Hippotragus leucophaeus*, MW222233, Hempel et al. 2021a). Only merged reads were used. "Shifted" refers to the second mapping to a reference with 200 bp shifted from the end to the beginning of the reference. The paired-end run of NBC RB4 D3 was split by the Sequence Read Archive into four files.

see supplementary file S2

**Supplementary table S4.** Paired-end sequencing results for the roan (*Hippotragus equinus*, Gonçalves et al. 2021) and sable antelope (*Hippotragus niger*, Koepfli et al. 2019) and the scimitar-horned oryx (*Oryx dammah*, Humble et al. 2020), mapped to the nuclear genome assembly of the scimitar-horned oryx (https://www.dnazoo.org/assemblies/Oryx_dammah, Humble et al. 2020). Merged and unmerged reads were used.

see supplementary file S2

**Supplementary table S5.** Paired-end sequencing results for the roan (*Hippotragus equinus*) (Gonçalves et al. 2021) and sable antelope (*Hippotragus niger*) (Koepfli et al. 2019) and the scimitar-horned oryx (*Oryx dammah*) (Humble et al. 2020) mapped to the mitochondrial genome assembly of the roan and sable antelope and the scimitar-horned oryx (JN632647, JN632648, JN632677; Hassanin et al. 2012). Merged and unmerged reads were used. "Shifted" refers to the second mapping to a reference with base pairs shifted from the end to the beginning of the reference (290bp for the scimitar-horned oryx, 252bp for the sable antelope and 300bp for the roan antelope reference).

see supplementary file S2

**Supplementary table S6.** Autosomal scaffold IDs larger than 1 Mb with scaffold size in parentheses of the scimitar-horned oryx (*Oryx dammah*) used in this study. Autosomal scaffolds were used as determined in Hempel et al. (2021b) by aligning the genome of the scimitar-horned oryx (Humble et al. 2020) to the X chromosome of the domestic goat (CM001739.2, Dong et al. 2013), the Y chromosome of the wild goat (CM003213.1, Dong et al. 2015) and the mitochondrial genome of the scimitar-horned oryx (JN632677, Hassanin et al. 2012) using SatsumaSynteny v2.0 (Grabherr et al. 2010).

| **Scaffold IDs larger 1 Mb of scimitar-horned oryx** | |
|---|---|
| HiC_scaffold_1 (111,031,402 bp),<br>HiC_scaffold_4 (90,666,804 bp),<br>HiC_scaffold_5 (107,200,824 bp),<br>HiC_scaffold_8 (61,296,289 bp),<br>HiC_scaffold_9 (60,415,816 bp),<br>HiC_scaffold_10 (82,229,679 bp),<br>HiC_scaffold_11 (51,053,030 bp),<br>HiC_scaffold_13 (64,050,454 bp),<br>HiC_scaffold_14 (50,506,852 bp),<br>HiC_scaffold_16 (67,964,418 bp),<br>HiC_scaffold_20 (71,114,058 bp),<br>HiC_scaffold_25 (105,560,413 bp),<br>HiC_scaffold_26 (100,398,400 bp),<br>HiC_scaffold_28 (118,425,367 bp),<br>HiC_scaffold_11045 (1,665,326 bp), | HiC_scaffold_11050 (12,440,347 bp),<br>HiC_scaffold_11056 (3,214,039 bp),<br>HiC_scaffold_11058 (2,359,909 bp),<br>HiC_scaffold_11072 (3,672,348 bp),<br>HiC_scaffold_11077 (8,331,692 bp),<br>HiC_scaffold_11081 (3,557,701 bp),<br>HiC_scaffold_11151 (1,109,432 bp),<br>HiC_scaffold_11163 (6,504,183 bp),<br>HiC_scaffold_11164 (7,448,190 bp),<br>HiC_scaffold_11203 (7,326,413 bp),<br>HiC_scaffold_11209 (5,441,144 bp),<br>HiC_scaffold_11210 (8,020,263 bp),<br>HiC_scaffold_11227 (4,755,615 bp),<br>HiC_scaffold_11229 (6,595,699 bp),<br>HiC_scaffold_11232 (7,227,722 bp) |

**Supplementary table S7.** *D* values, Z-scores, number of ABBA and BABA sites, and weighted block jacknife *D* values from computing *D* statistics using Dstat (transversion only version, https://github.com/jacahill/Admixture) with individuals in the following arrangement: P1: sable antelope (SB2152, Koepfli et al. 2019), P2: blue antelope, P3: roan antelope (10954, Gonçalves et al. 2021), P4: scimitar-horned oryx (SB20612, Humble et al. 2020).

| **Blue antelope specimen** | ***D* value** | **# ABBA sites** | **# BABA sites** | **Weighted block jackknife D value** | **Z-score** |
|---|---|---|---|---|---|
| NBC RB4 D3 | 0.1444 | 66054 | 49380 | 0.0073 | 19.7664 |
| NRM 590107 | 0.1240 | 75996 | 59227 | 0.0070 | 17.6837 |

**Supplementary table S8.** Number of trees found for sliding window tree analyses using WindowTrees v1.0.0 (https://github.com/achimklittich/WindowTrees) to determine gene flow with a window size of 100 kb and a threshold for missing data of 50% in binary mode with the individuals of blue antelope (NBC RB4 D3, NRM 590107, both this study), sable antelope (SB2152, HN216; Koepfli et al. 2019) and roan antelope (10954, He108; Gonçalves et al. 2021). The scimitar-horned oryx was used as outgroup (SB20612, Humble et al. 2020).

| | |
|---|---|
| **10954, He108, SB2152, NBC RB4 D3** | |
| (((SB2152,NBCRB4D3),(10954,He108)),SB20612) | 7,003 |
| (((NBCRB4D3,(10954,He108)),SB2152),SB20612) | 2,705 |
| ((NBCRB4D3,((10954,He108),SB2152)),SB20612) | 1,115 |
| (((He108,(SB2152,NBCRB4D3)),10954),SB20612) | 12 |
| (((SB2152,(He108,NBCRB4D3)),10954),SB20612) | 4 |
| sum | 10,839 |
| **10954, He108, SB2152, NRM 590107** | |
| (((SB2152,NRM590107),(10954,He108)),SB20612) | 6,802 |
| (((NRM590107,(10954,He108)),SB2152),SB20612) | 2,923 |
| ((NRM590107,((10954,He108),SB2152)),SB20612) | 1,068 |
| (((He108,(SB2152,NRM590107)),10954),SB20612) | 12 |
| (((SB2152,(He108,NRM590107)),10954),SB20612) | 4 |
| ((((NRM590107,He108),10954),SB2152),SB20612) | 1 |
| Sum | 10,810 |
| **10954, He108, HN216, NRM590107** | |
| (((HN216,NRM590107),(10954,He108)),SB20612) | 6,819 |
| (((NRM590107,(10954,He108)),HN216),SB20612) | 2,936 |
| ((NRM590107,((10954,He108),HN216)),SB20612) | 1,038 |
| (((He108,(HN216S,NRM590107)),10954),SB20612) | 11 |
| (((HN216,(He108,NRM590107)),10954),SB20612) | 4 |
| (((10954,(He108,NRM590107)),HN216),SB20612) | 1 |
| sum | 10,809 |

| **10954, He108, HN216, NBCRB4D3** | |
|---|---|
| (((HN216,NBCRB4D3),(10954,He108)),SB20612) | 6,976 |
| (((NBCRB4D3,(10954,He108)),HN216),SB20612) | 2,770 |
| ((NBCRB4D3,((10954,He108),HN216)),SB20612) | 1,076 |
| (((He108,(HN216,NBCRB4D3)),10954),SB20612) | 10 |
| (((HN216,(He108,NBCRB4D3)),10954),SB20612) | 5 |
| Sum | 10,837 |

**Supplementary table S9.** Number of tree topologies found for the sliding window tree analysis to determine gene flow directionality using WindowTrees v1.0.0 (https://github.com/achimklittich/WindowTrees) with a window size of 20 kb, a gap size of 80 kb between windows and a missing data threshold of 50% in binary mode with one individual per species: roan antelope (10954, Gonçalves et al. 2021), blue antelope (NBC RB4 D3, this study), sable antelope (SB2152, Koepfli et al. 2019) and the scimitar-horned oryx (SB20612, Humble et al. 2020) as outgroup.

| **Tree** | **Topology** | **# topologies found** |
|---|---|---|
| 1 | (((NBCRB4D3,10954),SB2152),SB20612) | 3,209 |
| 2 | (((NBCRB4D3,SB2152),10954),SB20612) | 5,437 |
| 3 | ((NBCRB4D3,(SB2152,10954)),SB20612) | 2,128 |

**Supplementary table S10.** Matrix of differing alleles for all pairwise comparisons for the eight sable antelope (*Hippotragus niger*) individuals using raw (below the diagonal)/normalised data (above the diagonal, data normalised by dividing by the number of sites used in the analysis - 2,243,953 sites) to estimate the species-wide nuclear diversity in relation to heterozygosity and to compare it to the other species. Normalised values plotted in fig. 5. Data used are from Koepfli et al. (2019).

| | **HN216** | **HN250** | **SB134** | **SB381** | **SB1954** | **SB2027** | **SB2130** | **SB2152** |
|---|---|---|---|---|---|---|---|---|
| **HN216** | x | 0.01283 | 0.01272 | 0.01406 | 0.01265 | 0.01222 | 0.0127 | 0.01283 |
| **HN250** | 28,784 | x | 0.01028 | 0.01160 | 0.01018 | 0.01355 | 0.01016 | 0.01032 |
| **SB134** | 28,539 | 23,060 | x | 0.01126 | 0.00858 | 0.01352 | 0.00950 | 0.00962 |
| **SB381** | 31,545 | 26,035 | 25,277 | x | 0.01106 | 0.01477 | 0.01087 | 0.01107 |
| **SB1954** | 28,396 | 22,838 | 19,262 | 24,822 | x | 0.01342 | 0.00967 | 0.00977 |
| **SB2027** | 27,428 | 30,409 | 30,344 | 33,132 | 30,118 | x | 0.01344 | 0.01355 |
| **SB2130** | 28,491 | 22,801 | 21,323 | 24,388 | 21,702 | 30,167 | x | 0.00977 |
| **SB2152** | 28,792 | 23,151 | 21,594 | 24,839 | 21,931 | 30,407 | 21,917 | x |

**Supplementary table S11.** Matrix of differing alleles for all pairwise comparisons for the three roan antelope (*Hippotragus equinus*) individuals using raw (below the diagonal)/normalised data (above the diagonal, data normalised by dividing by the number of sites used in the analysis - 2,243,953 sites) to estimate the species-wide nuclear diversity in relation to heterozygosity and to compare it to the other species. Normalised values plotted in fig. 5. Data used are from Gonçalves et al. (2021).

| | **He95** | **He108** | **10954** |
|---|---|---|---|
| **He95** | x | 0.01558 | 0.01500 |
| **He108** | 34,958 | x | 0.01595 |
| **10954** | 33,664 | 35,791 | x |

**Supplementary table S12.** Matrix of differing alleles for all pairwise comparisons for the two blue antelope (*Hippotragus leucophaeus*) individuals using raw (below the diagonal)/normalised data (above the diagonal, data normalised by dividing by the number of sites used in the analysis - 2,243,953 sites) to estimate species-wide nuclear diversity and to compare it to the other species. Normalised value plotted in fig. 5.

| | NRM 590107 | NBC RB4 D3 |
|---|---|---|
| **NRM 590107** | x | 0.00296 |
| **NBC RB4 D3** | 6,649 | x |

**Supplementary table S13.** Matrix of differing alleles for all pairwise comparisons for the eight sable antelope (*Hippotragus niger*) individuals using raw (below the diagonal)/normalised data (above the diagonal, data normalised by dividing by the number of sites used in the analysis - 4,596 sites) for genomes subsampled to 2.14x mean coverage to exclude the possible effect of sequence coverage on the species-wide nuclear diversity estimate. Data used are from Koepfli et al. (2019).

| | HN216 | HN250 | SB134 | SB381 | SB1954 | SB2027 | SB2130 | SB2152 |
|---|---|---|---|---|---|---|---|---|
| **HN216** | x | 0.01001 | 0.01001 | 0.01044 | 0.01023 | 0.00936 | 0.00914 | 0.01066 |
| **HN250** | 46 | x | 0.00979 | 0.01001 | 0.00827 | 0.01284 | 0.00870 | 0.00936 |
| **SB134** | 46 | 45 | x | 0.00783 | 0.00587 | 0.0111 | 0.00696 | 0.00675 |
| **SB381** | 48 | 46 | 36 | x | 0.00805 | 0.01197 | 0.00783 | 0.00849 |
| **SB1954** | 47 | 38 | 27 | 37 | x | 0.01088 | 0.00762 | 0.0074 |
| **SB2027** | 43 | 59 | 51 | 55 | 50 | x | 0.01066 | 0.01088 |
| **SB2130** | 42 | 40 | 32 | 36 | 35 | 49 | x | 0.00762 |
| **SB2152** | 49 | 43 | 31 | 39 | 34 | 50 | 35 | x |

**Supplementary table S14.** Matrix of differing alleles for all pairwise comparisons for the three roan antelope (*Hippotragus equinus*) individuals using raw (below the diagonal)/normalised data (above the diagonal, data normalised by dividing by the number of sites used in the analysis - 4,596 sites) for genomes subsampled to 2.14x mean coverage to exclude the possible effect of sequence coverage on the species-wide nuclear diversity estimate. Data used are from Gonçalves et al. (2021).

| | **He95** | **He108** | **10954** |
|---|---|---|---|
| **He95** | x | 0.0148 | 0.01523 |
| **He108** | 68 | x | 0.01436 |
| **10954** | 70 | 66 | x |

**Supplementary table S15.** Matrix of differing alleles for all pairwise comparisons for the two blue antelope (*Hippotragus leucophaeus*) individuals using raw (below the diagonal)/normalised data (above the diagonal, data normalised by dividing by the number of sites used in the analysis - 4,596 sites) for genomes subsampled to 2.14x mean coverage to exclude the possible effect of sequence coverage for the species-wide nuclear diversity estimate.

| | **NRM 590107** | **NBC RB4 D3** |
|---|---|---|
| **NRM 590107** | x | 0.00283 |
| **NBC RB4 D3** | 13 | x |

**Supplementary table S16.** Estimated autosomal heterozygosity for all scaffolds larger 1 Mb computed in ANGSD v0.923 (Korneliussen et al. 2014) for all roan and sable antelope individuals using 1) non-subsampled data, or 2) data subsampled once to that of the individual of each species with the lowest mean coverage (roan: 7.18x, sable antelope: 4.26x), and 3) data subsampled once to the level of the roan and sable antelope with the lowest mean coverage in the roan/sable antelope data set (4.26x). Data for roan and sable antelope from Gonçalves et al. (2021) and Koepfli et al. (2019), respectively.

| Sample ID | Species | Heterozygosity | | |
|---|---|---|---|---|
| | | 1) Not subsampled (mean coverage) | 2) Subsampled to species level (mean coverage) | 3) Subsampled to 4.26x |
| 10954 | Roan antelope | 0.00191 (24.42x) | 0.00179 (7.18x) | 0.00176 |
| He95 | | 0.00173 (7.18x) | 0.00173 (7.18x) | 0.00167 |
| He108 | | 0.00192 (7.40x) | 0.00192 (7.18x) | 0.00189 |
| SB2152 | Sable antelope | 0.00152 (24.23x) | 0.00129 (4.26x) | 0.00129 |
| SB134 | | 0.00117 (4.58x) | 0.00115 (4.26x) | 0.00115 |
| HN216 | | 0.00171 (5.87x) | 0.00167 (4.26x) | 0.00167 |
| HN250 | | 0.00121 (6.78x) | 0.00115 (4.26x) | 0.00115 |
| SB381 | | 0.00154 (4.64x) | 0.00152 (4.26x) | 0.00152 |
| SB1954 | | 0.00121 (4.26x) | 0.00121 (4.26x) | 0.00121 |
| SB2027 | | 0.00111 (4.40x) | 0.0011 (4.26x) | 0.0011 |
| SB2130 | | 0.00124 (4.70x) | 0.00121 (4.26x) | 0.00121 |

**Supplementary table S17.** Mitochondrial sequences from GenBank included in the haplotype network built with POPART v1.7 (Bandelt et al. 1999; Leigh and Bryant 2015).

| **Accession number** | **Sample ID** | **Included in network with … mitochondrial genomes** | **Publication** |
|---|---|---|---|
| xxxxxxxxxx | NBC RB4 D3 | Complete & partial | this study |
| MW222233 | NRM 590107 | Complete & partial | Hempel et al. 2021a |
| MW222234 | UPS ZMC 78488 | Complete & partial | Hempel et al. 2021a |
| MW228401 | NMW ST 715 | Partial | Hempel et al. 2021a |
| MW228402 | RMNH.MAM.20681.a | Partial | Hempel et al. 2021a |

**Supplementary table S18.** Sample information for the measured fossil blue (*Hippotragus leucophaeus*) and roan antelope (*Hippotragus equinus*) lower fourth deciduous premolars housed at the Archaeology Unit, Iziko Museums of South Africa, Cape Town, South Africa. The sample from which DNA was extracted successfully is printed in blue.

| **Site** | **Specimen number** | **Species identification** | ***Side*** |
|---|---|---|---|
| Die Kelders Cave 1 | DK1 MJA/H6 | *Hippotragus leucophaeus* | Right |
| | DK1 6084 | *Hippotragus leucophaeus* | Left |
| | DK1 - | *Hippotragus leucophaeus* | Right |
| Blombos Cave | BBC G5b 99.3 | *Hippotragus leucophaeus* | Left |
| Boomplaas Cave | BPA O14/OCH3 | *Hippotragus leucophaeus* | Left |
| Nelson Bay Cave | NBC YSL5 B8 | *Hippotragus leucophaeus* | Left |
| | NBC YSL3 A3 | *Hippotragus leucophaeus* | Right |
| | NBC DS19 1 | *Hippotragus leucophaeus* | Left |
| | NBC RB4 D3 | *Hippotragus leucophaeus* | Left |
| | NBC YGL34 A7 | *Hippotragus leucophaeus* | Right |
| | NBC GBSL23 D6 | *Hippotragus leucophaeus* | Left |
| Klasies River Mouth 1 | KRM1 2050/1/14/ASP | *Hippotragus leucophaeus* | Right |
| | KRM1 2051/1/14/A | *Hippotragus leucophaeus* | Left |
| | KRM1 ? | *Hippotragus leucophaeus* | Left |
| | KRM1 20660 | *Hippotragus leucophaeus* | Left |
| | KRM1 23856 | *Hippotragus leucophaeus* | Left |
| | KRM1 2022 | *Hippotragus leucophaeus* | Left |
| | KRM1 23855 | *Hippotragus leucophaeus* | Right |
| | KRM1 200 | *Hippotragus leucophaeus* | Right |
| | KRM1 201 | *Hippotragus leucophaeus* | Left |
| | KRM1 23050 | *Hippotragus leucophaeus* | Left |
| | KRM1 ? | *Hippotragus leucophaeus* | Right |
| | KRM1 2010 | *Hippotragus leucophaeus* | Left |
| | KRM1 2011 | *Hippotragus leucophaeus* | Left |

| | | | |
|---|---|---|---|
| Boomplaas Cave | BPA BRL6A P13 | *Hippotragus equinus* | Right |
| Nelson Bay Cave | NBC BSJ4 B1 | *Hippotragus equinus* | Right |
| | NBC CS16 D5 | *Hippotragus equinus* | Right |
| | NBC CS23 D3 | *Hippotragus equinus* | Right |
| | NBC BSJ105 B7 | *Hippotragus equinus* | Right |
